# Evolution of cooperation in multichannel games on multiplex networks

**DOI:** 10.1101/2024.09.19.613863

**Authors:** Amit Basak, Supratim Sengupta

## Abstract

Humans navigate diverse social relationships and concurrently interact across multiple social contexts. An individual’s behavior in one context can influence behavior in other contexts. Different payoffs associated with interactions in the different domains have motivated recent studies of the evolution of cooperation through the analysis of multichannel games where each individual is simultaneously engaged in multiple repeated games. However, previous investigations have ignored the potential role of network structure in each domain and the effect of playing against distinct interacting partners in different domains. Multiplex networks provide a useful framework to represent social interactions between the same set of agents across different social contexts. We investigate the role of multiplex network structure and strategy linking in multichannel games on the spread of cooperative behavior in all layers of the multiplex. We find that multiplex structure along with strategy linking enhances the cooperation rate in all layers of the multiplex compared to a well-mixed population, provided the network structure is identical across layers. The effectiveness of strategy linking in enhancing cooperation depends on the degree of similarity of the network structure across the layers and perception errors due to imperfect memory. Higher cooperation rates are achieved when the degree of structural overlap of the different layers is sufficiently large, and the probability of perception error is relatively low. Our work reveals how the social network structure in different layers of a multiplex can affect the spread of cooperation by limiting the ability of individuals to link strategies across different social domains.

## 1. Introduction

The conundrum of cooperation stems from the fact that altruistic behaviour involves a cost while selfish behaviour does not. Despite this disadvantage, cooperation is observed across all biological scales and several mechanisms have been proposed [1, 2, 3] to explain how altruistic behaviour can be sustained in a population. The Prisoners’ Dilemma (PD) game and its variants is often used as a prototypical model to understand evolution of cooperation in well-mixed as well as structured populations since it highlights the tension between shortterm personal gain and benefits of mutual cooperation. Two of those mechanisms that are of of significant importance in the context of our work are direct reciprocity [1, 4, 5, 6, 7, 8, 9, 10, 11, 12, 13, 14, 15, 16, 17, 18, 19, 20, 21, 22] and network reciprocity [23, 24, 25, 26, 27, 28, 29, 30, 31]. The success of direct reciprocity as a strategy for enhancing cooperation is contingent on repeated interactions between a pair of players.

In such a scenario, individuals can use conditional strategies [5, 6, 7] to retaliate against selfish behavior in future rounds. Previous studies have identified successful strategies in repeated games [10, 13, 14, 15, 16, 9, 17] and the conditions under which they can evolve [18, 19, 32, 20, 22, 21].

Network reciprocity also plays a significant role in enhancing cooperation levels. In network structured population, interactions of agents are constrained to their immediate network neighborhood. The formation of clusters of cooperators [25, 26] can enable members of the cluster to reap the benefits of mutual cooperation and prevent defectors from overexploiting them. The spread of cooperation then depends on the benefit-to-cost ratio of cooperative behaviour which is crucial in determining whether clusters of cooperators are likely to grow or shrink [23, 24, 25, 26, 27, 28, 29, 30, 31]. Simple rules for the evolution of cooperation under different network structures and different update rules, in large [26] and finite [31] populations have been uncovered for PD games.

Certain studies have integrated these two key mechanisms[33, 34, 35, 36, 37] of direct reciprocity and network reciprocity in order to understand their impact on the evolution of cooperation. These studies highlight the role of assortative interactions [34, 36], nature of the update process and network connectivity [35] in facilitating the evolution of cooperation. However, most existing studies predominantly focus on a single network structured population, and fail to capture the complexity of human interactions that occur across multiple domains. In reality, people and even organizations frequently engage with others in multiple social contexts concurrently and behaviour in one context can influence decisions in other contexts. For instance, academics can socially encounter colleagues in the workplace some of whom also happen to be their collaborators in different research projects. Individuals can belong to different online social networks and different companies are often found to compete in different geographical locations [38]. In each of these cases, regardless of whether the nodes of the network represent individuals or organizations, interaction partners across different domains can either be the same or distinct for each domain. Competition and conflicts in one domain can overtly or subtly affect interactions in the other domains [39].

Despite the complex and unpredictable nature of social environments, humans excel in processing social information and can tailor their behaviour accordingly [40]. The benefits yielded and costs incurred by altruistic behaviour can vary across social domains. As a result, people tend to behave differently in domains with different interacting neighbors and can also couple their behaviours across different social domains while dealing with interacting partners that are common across those domains. It is therefore imperative to move beyond well-mixed populations and single layer networks in order to understand the impact of multi-level interactions on the evolution of cooperation. Multiplex networks provide the ideal framework [41, 42, 43, 40] for addressing these issues [44, 45, 46, 47, 48, 49, 50, 51, 52]. The different layers of a 2-layer multiplex network have been used [53, 54] to distinguish between the social network where the game is played (called the interaction network) and the strategy update network that is used to update strategy through imitation [46, 52]. Several groups have analysed evolutionary games on multiplex structured networks and highlighted the benefits afforded by multiple layers [45, 48], asynchronous strategy update [50] and structural similarity between layers [49] in promoting cooperation in such systems. In all these models, the different layers of the multiplex were coupled through total payoff (calculated by adding the payoffs from interactions with all neighbours across all layers of the multiplex) and the strategy update process (where an individual in one layer can update her strategy by imitating the strategy of a neighbour in a different layer). They however disregarded the interplay between an individual’s behaviour in one layer and its potential influence in another layer.

A recent study by Donahue *et al*. [55] have highlighted the positive impact of linking strategies across multiple games on the evolution of cooperation. They found that coordinating strategies across multiple games can lead to enhanced cooperation levels across all games. However, such investigations have ignored the potential role of network structure, focusing only on analysing the effects of linking strategies across multiple games occurring in a single well-mixed population.

In this paper, we use a multiplex network to describe social interactions that occur across multiple domains. A population of individuals simultaneously engage in multiple *repeated* PD games, each occurring on a different layer of the multiplex. The different games are distinguished by distinct benefit-to-cost ratio for altruistic behaviour. The network structure on each layer of the multiplex can either be identical to or distinct from other layers with the deviation from complete structural overlap of the layers being determined by the average edge overlap 𝒪between the two layers of the multiplex. As a consequence, each individual can have a set of connections with neighbours in one layer that are distinct from her connections in another layer (0 ≤𝒪 *<* 1). When there exists complete structural overlap between the layers (𝒪 = 1), each individual has the same set of connections across all layers of the multiplex. Individuals can link their strategies from multiple games against their common interaction partners across layers (represented by black edges in Fig. 1a) to induce cooperative behaviours in those social contexts where cooperation is relatively costly. However, an agent treats each game independently while playing against unique interaction partners (represented by red edges in Fig. 1a) in different layers. Each individual’s payoffs arising from interactions in each layer are aggregated across all layers. In our model, layers of multiplex are not only coupled through payoffs; individuals can also link their behaviour against common neighbors across layers. By studying repeated PD games on a multiplex, we wish to investigate the impact of multiplex structure, degree of structural overlap between layers, variable benefits of cooperation across different layers on the emergent levels of cooperation. Our model extends beyond analysing the role of multiplex network structures in multichannel games. Using a theoretical framework, we also explore the effect of perception errors in recalling past actions, due to imperfect memory, in multiple repeated games with both unlinked and linked strategies.

**Figure 1:**
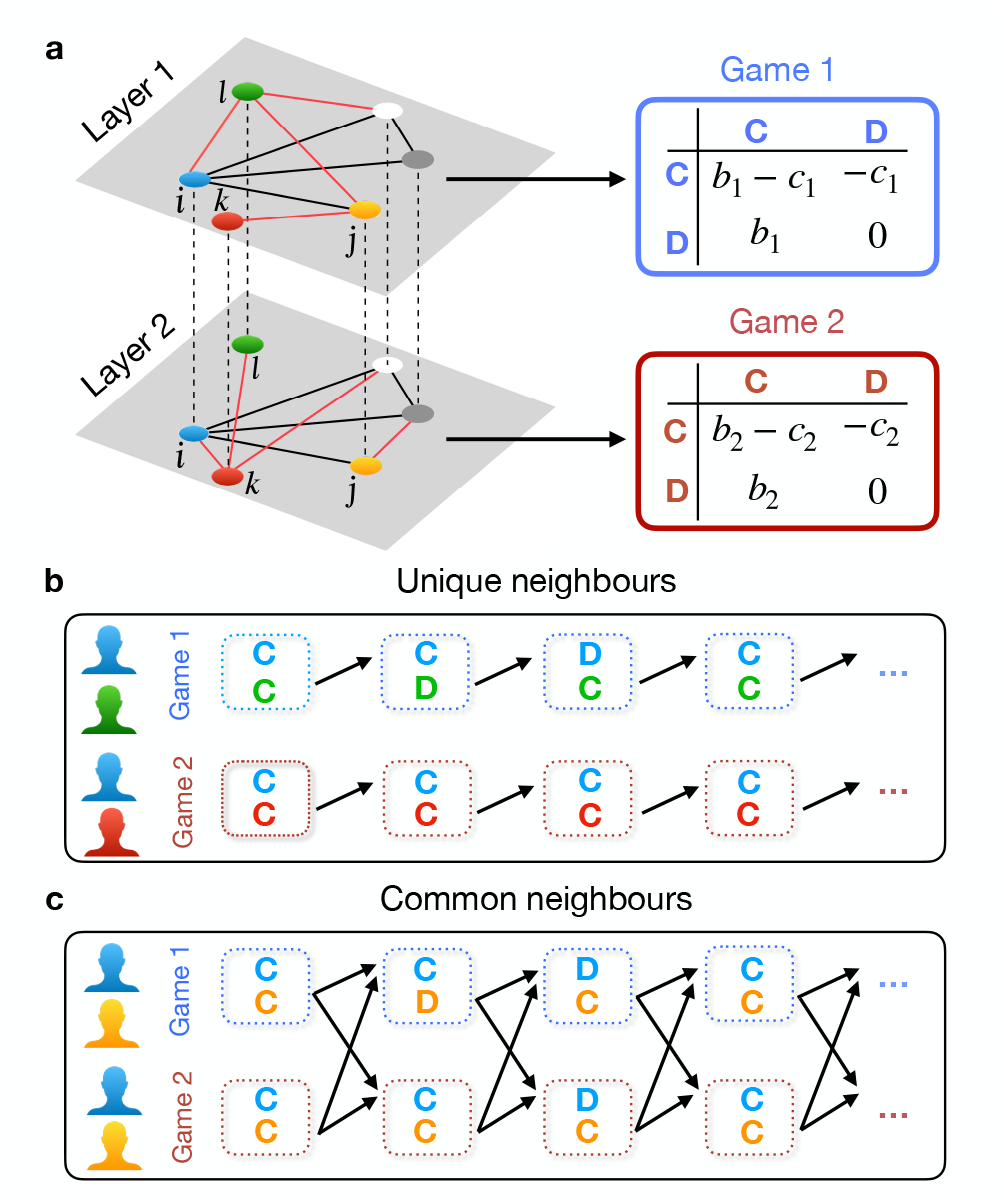
Schematic of multichannel games on a two-layer multiplex network: **(a)** Two distinct donation games, each with different benefits and costs of cooperation, are played across the layers. Each individual is represented by a node in layer one, with dotted lines indicating connections to their counterparts in the other layer. Red edges denote unique connections, while black edges signify common connections between pairs of nodes across both layers. **(b)** Players treat each game independently when interacting with unique neighbors in each layer. A player (blue) responds to a unique neighbor’s (green in layer 1, red in layer 2) previous action within the same game and layer. This is indicated by an arrow from the opponent’s action in the previous round to the focal player’s response (action) in the current round. **(c)** When interacting with a common partner, a player (blue) can link the two games, reacting to the opponent’s (yellow) defection in one game by defecting in both games. This linkage is assumed to occur only in interactions with common neighbors. This is indicated by two arrows pointing from the common opponent’s actions in the last round in both games to the player’s actions in the current round.

Our results indicate that the outcome of evolutionary dynamics in multiplex network structured populations is often in stark contrast to that in well-mixed populations. This is observed both in the absence and presence of strategy linking. We find that strategy linking against common neighbors is only effective in promoting cooperation when the degree of edge overlap between layers is substantial. Our results underscore the importance of population structure and imperfect memory on the evolution of cooperation in multiple repeated games.

The paper is structured as follows: Sec. 2 elaborates on the methodologies employed for calculating payoff and cooperation rate in the context of multiple repeated PD games on a multiplex network. In section 3, we present all the main results of the paper and finally conclude with a discussion of the key results and possible future directions of research in Sec.4.

## 2. Methods

We consider a population of individuals engaging in distinct Prisoner’s Dilemma (PD) games in different layers of an *m*-layer multiplex network, with each layer having an identical number (*N*) of nodes. Each individual is represented by a node that can potentially have different neighbors in different layers. Each layer *α* of the multiplex is represented by an adjacency matrix 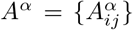, where 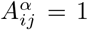 when individuals *i* and *j* are connected in layer *α*, and 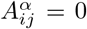 otherwise. A vector of adjacency matrices characterizes a multiplex network of *m* layers, **A** = {*A*^1^, *A*^2^, …, *A*^*m*^}.

### 2.1 Key properties of multiplex networks

The degree of a node *i* in a specific layer *α* of a multiplex is calculated as, 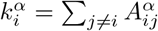 from which it follows that 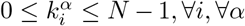. Thus, the multiplex degree of node *i* is a vector 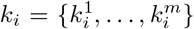. If, 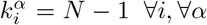, ∀*α*, then each layer of the multiplex network is a complete graph and the multiplex network reduces to a single well-mixed population.

A key feature of a multiplex network is that a pair of nodes can be connected in one layer but not in others. As illustrated in Fig. 1 which shows a multiplex network with two layers, a node may have a different set of neighbors in layer 2 compared to layer 1. For example, the node *j* acts as a common neighbor of node *i* in both layer 1 and layer 2 (indicated by black edges). However, node *i* is exclusively connected to node *k* in layer 2 (illustrated by a red edge), without a corresponding connection in layer 1.

To quantify the degree of similarity between layers, we consider the edge overlap between a pair of nodes *i* and *j*, defined as, 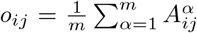, where *m* is the number of layers of a multiplex. *o*_*ij*_ is 1 if the edge *i*− *j* is common across all layers. The number of common neighbors 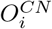 across a pair of layers *α, β* and unique neighbors 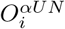 of a node *i* in layer *α* are respectively defined as

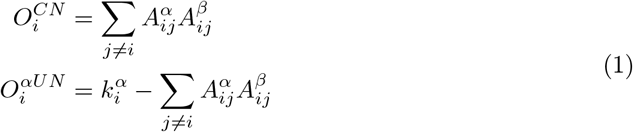

The average (avg.) fraction of common neighbors (i.e. edge overlap) across a pair of layers *α, β*, is given by:

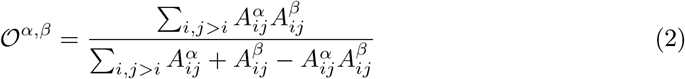

For all pairs of *m* layers of a multiplex[56], the above quantity can be generalized to 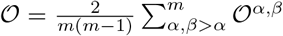. When all layers of a multiplex are the same (i.e. have identical topologies and edges), the average edge overlap of a multiplex becomes maximum, 𝒪= 1. Similarly, when all layers are completely distinct from each other (each node has different connections across layers), then edge overlap is minimum, 𝒪= 0. In our model the average fraction of *unique* neighbors is denoted by 𝒪−1. The algorithm used to create a multiplex network with desired edge-overlap between the layers is described in Sec.2 of the supplementary information (SI).

### 2.2 Strategy employed against common and unique neighbors

In our framework, two distinct PD games, characterized by distinct payoff matrices, are being played in the two layers of the multiplex network. Each individual engages in both games repeatedly and simultaneously with their respective neighbors in each layer, taking a decision to Cooperate (C) or Defect (D) in each round. Although the formalism described here is valid for any value of *m*, we have used *m* = 2 throughout this paper. Players use reactive strategies, where behavior depends only on the co-player’s previous actions. In our model, agents employ distinct strategies based on the nature (common or unique) of their neighbors across network layers. When interacting with common neighbors, the strategy is based on the collective actions of the co-player in the last round in *both* layers. Such strategies have been called [55] multi-game linked reactive strategies. Specifically, a player playing against a common neighbor (CN) in the two layers of a multiplex network, the player’s strategy takes the form of a 10 tuple,

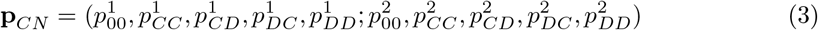

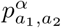 is the player’s probability to cooperate in layer *α*, depending on the co-player’s previous actions *a*_1_ and *a*_2_ in layer 1 and layer 2 respectively. The first five components of the strategy vector signify the player’s strategy in layer 1, while the last five components denote her strategy in layer 2. The first entry for each layer 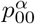 is the initial probability of cooperating against a common neighbor in layer *α*.

When playing against a unique neighbor (UN) in a specific layer *α*, players are unable to condition their behaviour on the opponent’s action in other layers, and strategy against such neighbors depends only on the co-player’s last-round action in the layer *α*. Such a strategy has been called [55] a multi-game unlinked reactive strategy and is defined as

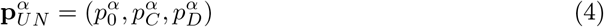

We note that the set of unique neighbor strategies is a strict subset of the common neighbor strategies (they correspond to those **p**_*CN*_ for which 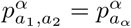 for all layer *α*, and all actions *a*_1_ and *a*_2_). We consider finitely repeated games with *w* as the probability of another round of games. For infinitely repeated prisoner’s dilemma game between the focal player and each of her neighbours in both layers, the first round probability to cooperate becomes irrelevant.

### 2.3 Payoff calculation

A player’s payoff in each game *α* (occurring in layer *α*) is either *R*_*α*_, *S*_*α*_, *T*_*α*_, or *P*_*α*_, depending on the player’s and the co-player’s action. We consider a donation game (a version of the PD game) for which payoff matrix elements become *R*_*α*_ = *b*_*α*_− *c*_*α*_, *S*_*α*_ = −*c*_*α*_, *T*_*α*_ = *b*_*α*_, and *P*_*α*_ = 0, where *b*_*α*_ and *c*_*α*_ are the benefit and cost of cooperation in game *α*.

The expected pairwise payoff for player *i* (or *j*) in the infinitely repeated game, when playing against a neighbor *j* (or *i*) in layer *α* are respectively given by

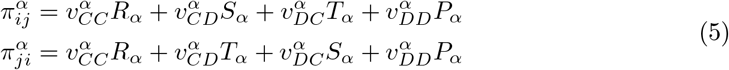

where 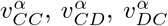, and 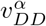 represent the equilibrium probability of finding the game in layer *α* in the state *aã* where *a,ã* correspond to the actions of player 1 and player 2 respectively in layer *α* (refer to Sec.3 of SI for detailed calculation of 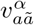).

The accumulated average payoff of player *i* in layer *α* due to pairwise interactions with all neighbors (including both common and unique neighbors) is,

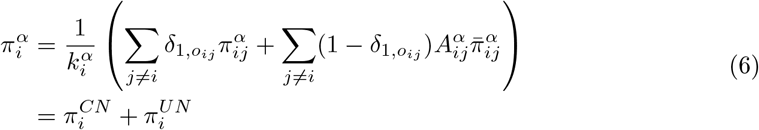

Where, the Kronecker delta term equals 1 when node *j* is a common neighbor of *i* across all layers (i.e. *o*_*ij*_ = 1) and is 0 otherwise. 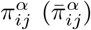) is pairwise payoff of *i* against a common (unique) neighbor *j* in layer *α* using strategy **p**_*CN*_ 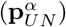 respectively.

The total expected payoff for player *i* as a consequence of her interactions with all neighbours across all layers of the multiplex network is

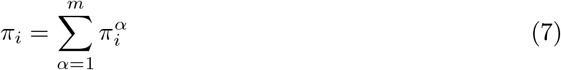

### 2.4 Cooperation rate calculation

Player *i*’s (or alternatively, player *j*’s) average cooperation rate when playing against player *j* (or alternatively, player *i*) in layer *α* respectively is,

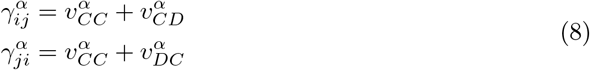

The average cooperation rate of player *i* against common and unique neighbors in layer *α* can be calculated separately by averaging her cooperation rate against each type of neighbor respectively, in that specific layer.

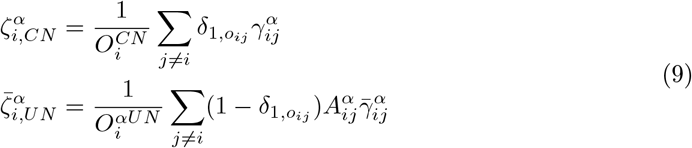

Where, 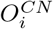 and 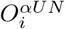 is number of common and unique neighbors (in layer *α*) respectively of player *i* and 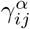 is cooperation rate of player *i* against common neighbor *j* in layer *α*. Similarly, 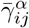 is cooperation rate of *i* against a unique neighbor *j* in layer *α*, calculated using the unlinked reactive strategy 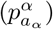. (For a detailed calculation of 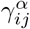 and 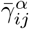 refer to Sec.3 of SI). Similarly, an individual *i*’s average cooperation rate in layer *α* against all of her neighbors is,

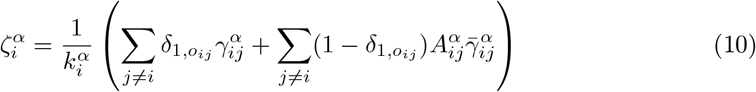

At each time step, the population’s average linked and unlinked cooperation rate in game *α* (occurring in layer *α*) against common neighbor and unique neighbors respectively is calculated by averaging over the whole population for each type separately.

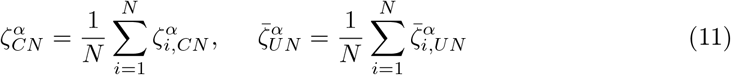

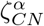 is population’s average linked cooperation rate against common neighbors, calculated using linked reactive strategies 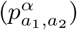 and similarly, 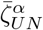 is population’s average unlinked cooperation rate against unique neighbors in layer *α*, calculated using unlinked reactive strategies 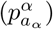.

At each time step, the population’s average cooperation rate is calculated as,

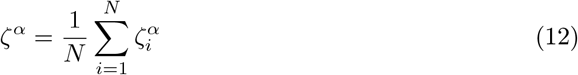

For all future instances, we will refer to the population’s average cooperation rate and average linked (unlinked) cooperation rate as cooperation rate and linked (unlinked) cooperation rate only.

### 2.5 Categorization of outcomes

Simulation outcomes are classified on the basis of the population’s cooperation rates [55].

Full cooperation: *ζ*^*α*^ ≥ 0.8 ∀ *α*

Only layer *α* cooperation: *ζ*^*α*^ ≥ 0.8 and *ζ*^*β*^ ≤ 0.2 ∀ *β* ≠ *α*

No cooperation: *ζ*^*α*^ ≤ 0.2 ∀ *α*

### 2.6 Strategy update

At each time step of the simulation, the population updates their strategy according to two different updating methods. They either explore random strategies within their respective strategy spaces with probability *µ*, or with probability 1− *µ*, imitate a randomly chosen neighbor’s strategy based on the neighbor’s perceived success [57, 58]. The imitation process followed the pairwise comparison rule [57, 58], where an individual *A*, would imitate the strategy of another individual, *B*, with a probability of *p* = (1 + exp(− *s*(*f*_*B*_− *f*_*A*_)))^−1^. Here, *f*_*A*_ (*f*_*B*_) represents the payoffs of individual *A*(*B*), and *s* denotes the intensity of selection. Since each individual is equipped with both multi-game linked and unlinked reactive strategies to interact with their common and unique neighbors respectively, both strategies of the focal player need to be updated. Consequently, during the imitation process, player *A* will adopt both strategies of player *B* according to the pairwise comparison rule. We have also explored two different update schemes.

#### i. Independent strategy update

At each time step, a single network layer was randomly selected, and all individuals within that network were given the opportunity to update their strategies.

#### ii. Simultaneous strategy update

At each time step, all individuals across all layers update their strategy. If a common neighbor is selected for imitation by the focal player in any one layer, the strategy of the focal player is updated across all layers. If an unique neighbor is selected for imitation in any one layer, the strategy of the focal player is updated only in the corresponding layer.

## 3. Results

We analyse the effect of complex network structure on the cooperation rate in multichannel games on multiplex networks by considering different types of complex networks, such as random-regular networks (RRN) and scale-free networks (SFN), having identical topologies and identical connections across all layers of the multiplex network. Such an analysis allows us to understand how the underlying network structure affects cooperation levels in contrast to well-mixed populations represented by a complete graph. We then analyse 2-layer multiplex networks having identical topologies but distinct connections across layers such that the same node of the network in different layers can have a different set of connections. This allows us to understand how the degree of edge overlap between identical nodes in the different layers affect cooperation levels in multichannel games where strategies can be linked only for interactions with those neighbours of a node that are common across both layers.

### 3.1 Evolutionary dynamics in multiplex networks having identical topologies and identical connections across layers

We work with a multiplex network having two layers and both layers have RRN topology with *N* nodes. Each node is connected at random to *k* other nodes. The *k* neighbours of each node are common across both layers of the multiplex network (𝒪 = 1) as a consequence of which the behaviour of each individual in the network can be linked across all layers of the multiplex. This enables each individual to condition her behaviour on the opponent’s actions in *all* games. For comparison, we also consider a scenario where individuals treat each game independently, conditioning their behavior on the opponent’s actions in that specific game. Since each node has the same set of connections in both layers, our model of a 2-layer network topology effectively reduces to a single network topology with maximal edge overlap. However, the two layers of the network can still be distinguished from each other if the two games played on the two networks are distinct, i.e. have different payoff matrices. This scenario generalizes the work of Donahue et al.[55], which considered a mixed population which arises as a limiting case of our model when networks in all layers are described by a complete graph i.e. a graph with *k* = *N*− 1.

We assume (unless specified otherwise) that the game played in layer 1 has a higher benefit of cooperation, such that *b*_1_ *> b*_2_. Despite the smaller benefit in the second game, very high cooperation evolves in both games, reaching 94.8% in the first game and 80.6% in the second, when individuals treat each game independently (Fig. 2a-d) and interact with just four neighbors (*k* = 4). As the degree of the multiplex network increases, high cooperation rate persists with unlinked strategies only in game 1; but the cooperation rate steadily decreases in game 2 (Fig. 2a). However, when individuals use linked strategies, higher cooperation rate is observed in *both games*, even for high degree (Fig. 2e). Figure 2b,f shows the temporal evolution of cooperation rates for unlinked and linked strategies, respectively, when each layer in the multiplex has a degree *k* = 4. In order to better understand these results, we plot the abundance of different evolved cooperation scenarios (see Sec. 2.5) in the population. Contrary to Donahue *et al*. [55], we find that the full cooperation scenario is the most likely evolutionary outcome for both unlinked (Fig. 2c) and linked (Fig. 2g) cases when individuals interact with only four neighbors, as opposed to a well-mixed population where everyone interacts with everyone else (*k* = *N* − 1). These results suggest that the multiplex network structure itself can promote cooperation in both layers when the connectivity of the network is low, irrespective of strategy linking. However, for high network connectivity (i.e. large *k*), linked strategies are more effective in promoting cooperation in both layers of the multiplex compared to unlinked strategies.

**Figure 2:**
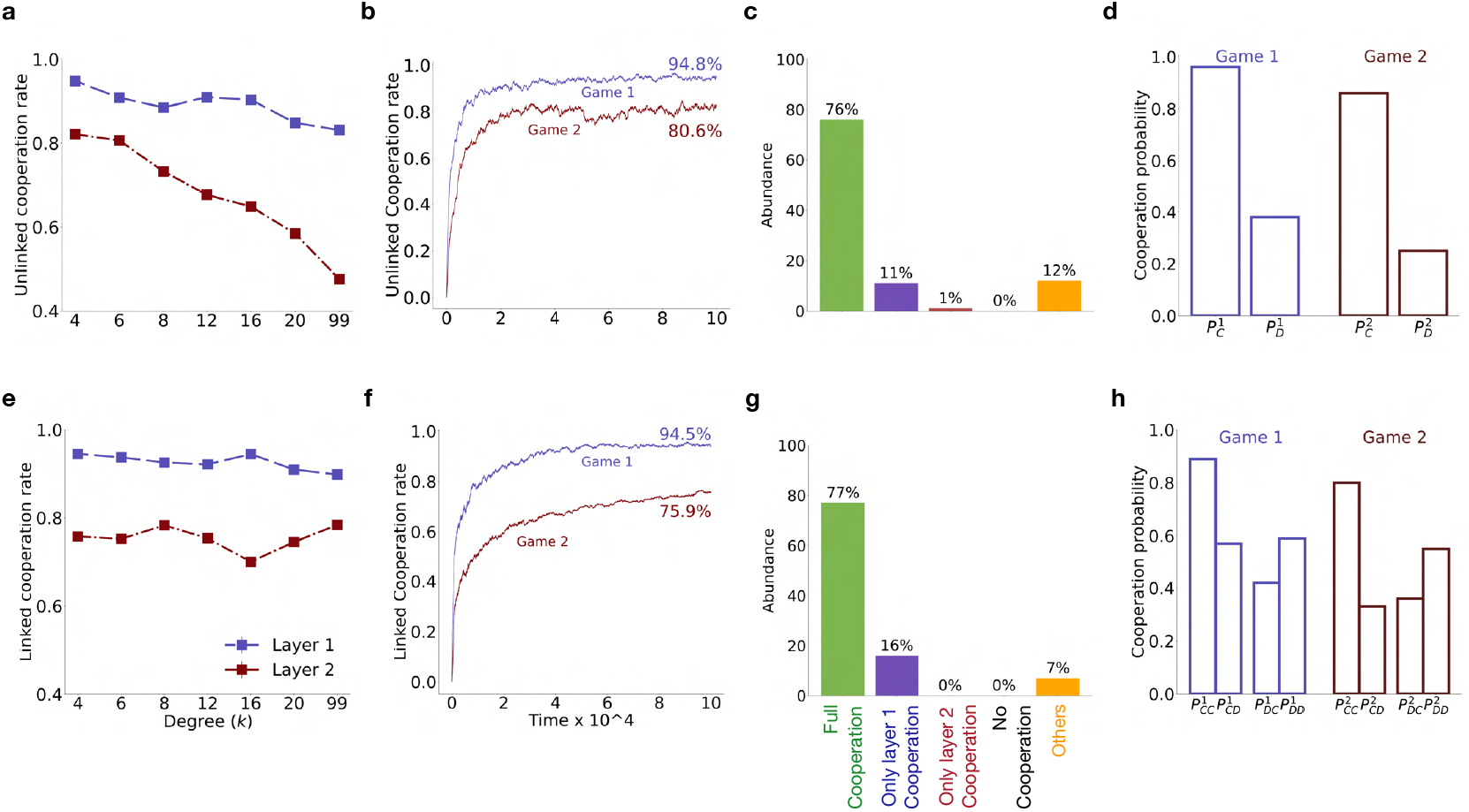
Impact of network connectivity. Evolution of cooperation in multiplex structured population using unlinked **(a-d)** and linked **(e-h)** strategies. **(a**,**e)** shows the variation of the cooperation rate with degree (*k*^1^ = *k*^2^ = *k*); **(b**,**f)** the time evolution of the cooperation rate for *k* = 4; **(c**,**g)** relative abundance of different cooperation scenarios across multiple simulations for *k* = 4; **(d**,**h)** the average strategy used for *k* = 4; when all individuals use unlinked and linked strategies respectively. The results were obtained by averaging over 200 simulations using the independent strategy update rule and RRN topology for each layer of the multiplex network. Other parameters used: *N* = 100, *b*_1_ = 5, *b*_2_ = 3, *c*_1_ = *c*_2_ = 1, *µ* = 0.001, *w* = 1, and *s* = 2.

To understand how a network structured population with degree *k* = 4 facilitate high cooperation in both games, we examine the population’s average strategy at the end of simulation. The average strategy in any layer *α* is obtained by averaging 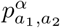 in the linked case (or 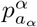 in the unlinked case) over the population and then further averaging the result over multiple trials. In the unlinked case (Fig. 2d), the average strategy in layer 1, resembles Generous Tit-for-Tat (GTFT), where individuals reciprocate altruistic behavior with a high probability 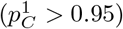 in layer 1 and still cooperate with a moderate probability 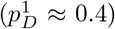 in response to a co-player’s defection. However, the evolved strategies in layer 2 are less cooperative 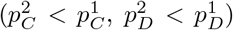, leading to a lower cooperation rate in game 2. In the linked case (Fig. 2h), on average, individuals cooperate with high probability across all layers in response to cooperation in both games, but lower their cooperation probability in both games if the co-player defects in either. Under the independent strategy update scheme, we observed that the evolved strategy in the population cooperates with high probability only when the co-player either cooperates or defects in both games in the previous round. For simulations using deterministic strategies, the evolved strategy can be represented as, **p**= (1, 0, 0, 1; 1, 0, 0, 1). This strategy ought to be vulnerable to invasion by ALLD strategy, which defects in both games. However, for an ALLD strategy to emerge, a single-layer defector strategy, e.g. **p**^*′*^= (0, 0, 0, 0; 1, 0, 0, 1) or (1, 0, 0, 1; 0, 0, 0, 0) has to first evolve in the resident population of the evolved cooperative strategy **p** under independent strategy update. But, when such a single-layer defector strategy starts defecting in one game, it is punished in both due to the low *p*_*CD*_ and *p*_*DC*_ values of the evolved strategy, **p** in both games. For an infinitely repeated multichannel game between **p** and **p**^*′*^, in the stationary distribution they will be in the *DDDC* or *DDCD* state of the Markov chain with a probability close to 1. As a result, the evolved cooperative strategy gets a payoff *b*_2_ or *b*_1_, whereas the single-layer defector strategy gets −*c*_2_ or −*c*_1_. Thus, the single-layer defector strategy is at a disadvantage in a population of strategies defined by **p**, making the emergence of the ALLD strategy very unlikely. Hence, evolved cooperative strategies such as **p** remain robust under the independent strategy update scheme.

When we simulated the evolutionary dynamics using the simultaneous strategy update scheme, a similar pattern was observed in the evolution of the linked cooperation rate in both layers (Fig. 3**a**) and the abundance of different cooperation scenarios (Fig. 3**b**). However, the average strategy in the full cooperation scenario (see Fig. 3**c**) are mutually cooperative with a high probability 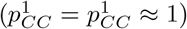 and tend to cooperate with a similar probability, comparable to what is expected by chance, when opponent defects in either or both games. In order to understand if the persistently higher linked cooperation rate is robust to changes in population size as well as network topology of the layers, we carried out simulations for networks of varying sizes as well as different topologies for two different update schemes and finitely repeated games in both layers. Our results generally indicate that a higher linked cooperation rate is observed in both Game 1 Fig.4a, 4d and Game 2 Fig.4b, 4e in structured populations (multiplex created with RRN and SFN), across all sizes, when compared with the mixed population scenario. The effect of network structure on the linked cooperation rate for both games is more pronounced for large population size when the simultaneous strategy-update scheme is used. As shown in Fig. 4c, 4f full cooperation scenario is more prevalent in structured populations as compared to mixed populations. We find that direct reciprocity along with strategy linking in multichannel games, even when acting together, may not be sufficient to ensure high levels of cooperation in well-mixed populations when the population size is large. However, the presence of a multiplex network structure in conjunction with strategy linking can significantly enhance cooperation in multichannel games (see Figs. 4d-4f) even for large populations.

**Figure 3:**
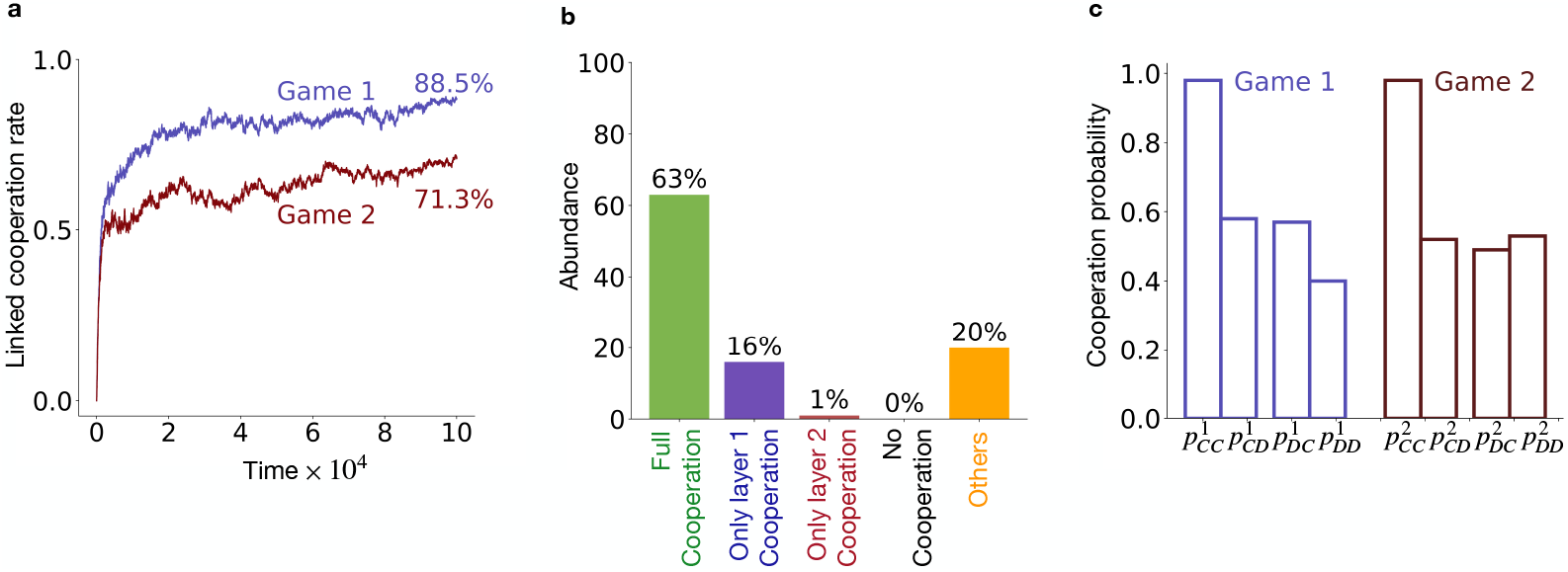
Evolutionary dynamics under the simultaneous strategy update scheme. **(a)** Evolution of linked cooperation rate in network layers 1 and 2. **(b)** Relative abundance of different evolved cooperation scenarios obtained across 100 independent simulations.**(c)** Population’s average strategy in the full cooperation scenario. Each bar represents population’s average of 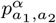 in the respective layer, averaged over 100 independent realisation where full cooperation scenarios evolved. The 2-layer multiplex network had identical RRN topology with *k* = 4 in each layer. All other parameter values are same as Fig.2.

**Figure 4:**
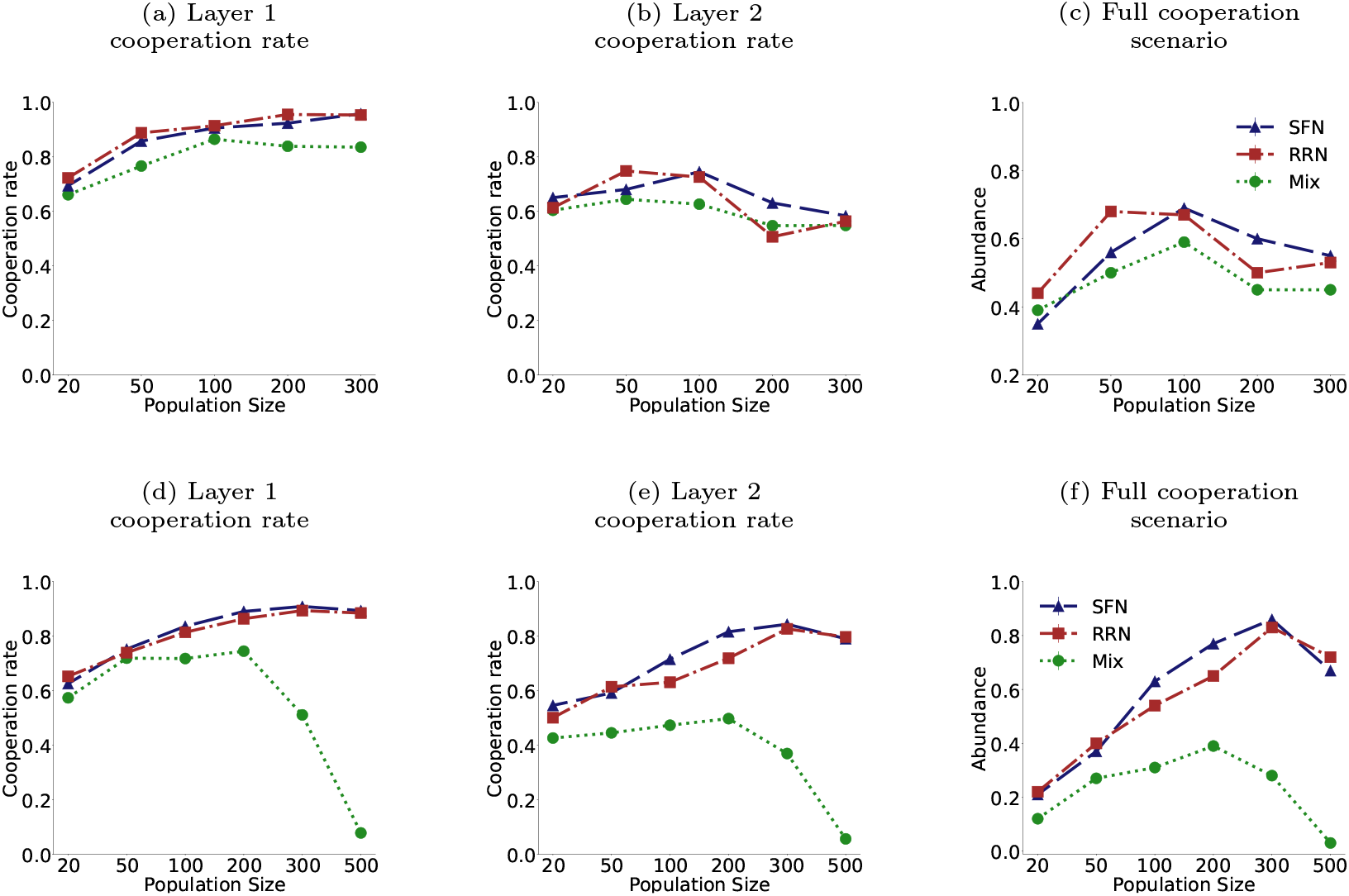
Comparison between evolved linked cooperation rates for different topologies and population sizes. Each data point depicts the cooperation rate of population in Layer 1 (**a**,**d**), in Layer 2 (**b**,**e**) at the end of the simulation and (**c**,**f**) the fractional abundance of full cooperation scenarios across 100 independent runs. Panels **a-c** correspond to the independent strategy update scheme and panels **d-f** correspond to simultaneous strategy update scheme respectively. *w* = 0.95 was used. For SFN networks, average degree 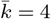. Other parameters values used are same as in Fig. 2

We have also verified that our results are in agreement with that of Donahue et al. [55] when the underlying network is a complete graph in both layers for the case where *N* = 50 and *µ* = 0.001 which corresponds to the low mutation limit as investigated in [55].

### 3.2 Evolutionary dynamics of a reduced set of strategies

We carried out an analysis using reduced strategy sets in order to better understand how the structure of the underlying multiplex network in multichannel games affects the nature of the emergent linked strategy.

For the purpose of our analysis, we consider a three-strategy system, consisting of Linked GTFT (LGTFT) = (1, 1, *q, q, q*; 1, 1, *q, q, q*), ALLD = (0,0,0,0,0; 0,0,0,0,0) and ALLC = (1,1,1,1,1; 1,1,1,1,1). The LGTFT strategy considered here is an approximation of the average strategy that evolved in the full cooperation scenario (*ζ*^*α*^≥ 0.8 ∀ *α*), for the simultaneous strategy update scheme (Fig. 3**c**). This strategy starts by cooperating and continues to cooperate in all games as long as the opponent also cooperates. However, if the opponent defects in any of the two games, LGTFT immediately lowers her cooperation probability in all games to 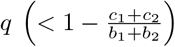. Due to the similarity of this strategy with the well-studied Generous Tit-for-Tat (GTFT) [6] that emerges in the context of single repeated games, we have named this strategy LGTFT. In the limit *q* = 0, the LGTFT strategy reduces to the Linked TFT (LTFT) strategy.

We first analyse a 2-strategy system to identify the conditions under which the LGTFT strategy can invade a population of ALLD players in both well-mixed and structured populations. The conditions, 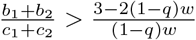 and 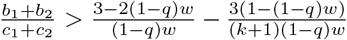 respectively, translate to lower bounds on the benefit (*b*_1_) in game 1 (see Sec.4.1 of SI for details). We then carry out simulations, in the absence of mutations, to obtain the fixation probability of the LGTFT strategy in a resident population of ALLD for different values of the benefit in game 1 (*b*_1_) while keeping the benefit in game 2 (*b*_2_) fixed (Fig. 5). In a *large but finite* population, we observe that the critical benefit required in game 1 for the fixation probability of LGTFT to exceed the neutral fixation probability of 1*/N* is significantly lower when interactions of each node are constrained to a few connected neighbors (for example, when *k* = 3, 5) compared to the well-mixed case (where *k* = *N* − 1) and agree well with our theoretical predictions (see Sec.4.1 of SI for details).

**Figure 5:**
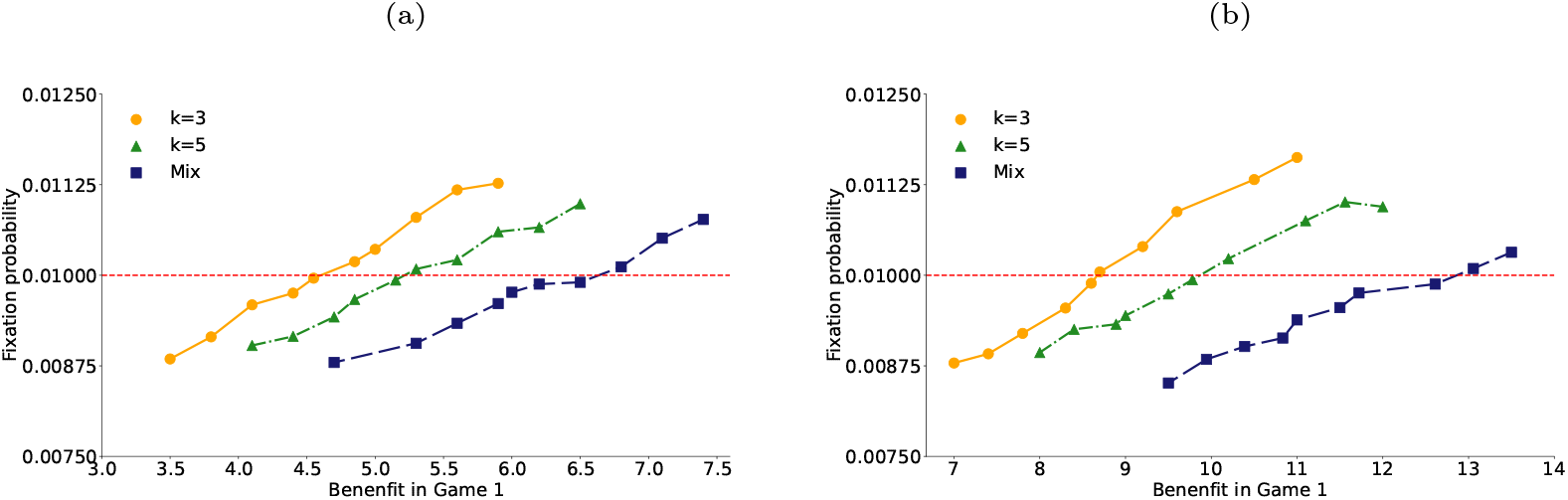
Variation of fixation probability of the LGTFT strategy with benefit in game 1 for different degrees. **(a)** *q* = 0, **(b)** *q* = 0.3. The fixation probability was calculated from 10^6^ trials. The dotted horizontal line shows the neutral fixation probability. The network was created using RRN topology. Other parameters used: *b*_2_ = 2, *c*_1_ = *c*_2_ = 1, *w* = 0.5, *s* = 0.01, *N* = 100.

In a system with three strategies, the abundance of both LGTFT and ALLD strategies exceeds 1*/*3 in both well-mixed and structured populations in the weak selection limit when 2*q*^*^ − 1 *< q < q*^*^, where 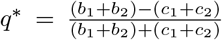. However, for LGTFT to dominate over both ALLD and ALLC, the conditions *q <* 2*q*^*^ − 1 and 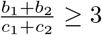 need to be simultaneously satisfied (see section 4.2 of SI for details). Numerical simulations of the competition between these three strategies provides additional insights into the dynamics for arbitrary selection strengths. Fig. 6 shows a heat map depicting the relative abundance of the three strategies in the low mutation limit. Each point in the simplex *S*_3_ represents the long-term average frequency of all three strategies in the population, with the vertices representing scenarios where only one of the three strategies are present in the population. The color-coding of each point in *S*_3_ represents the number of times, out of 1000 realizations, the population’s composition was represented by the corresponding point in the simplex.

**Figure 6:**
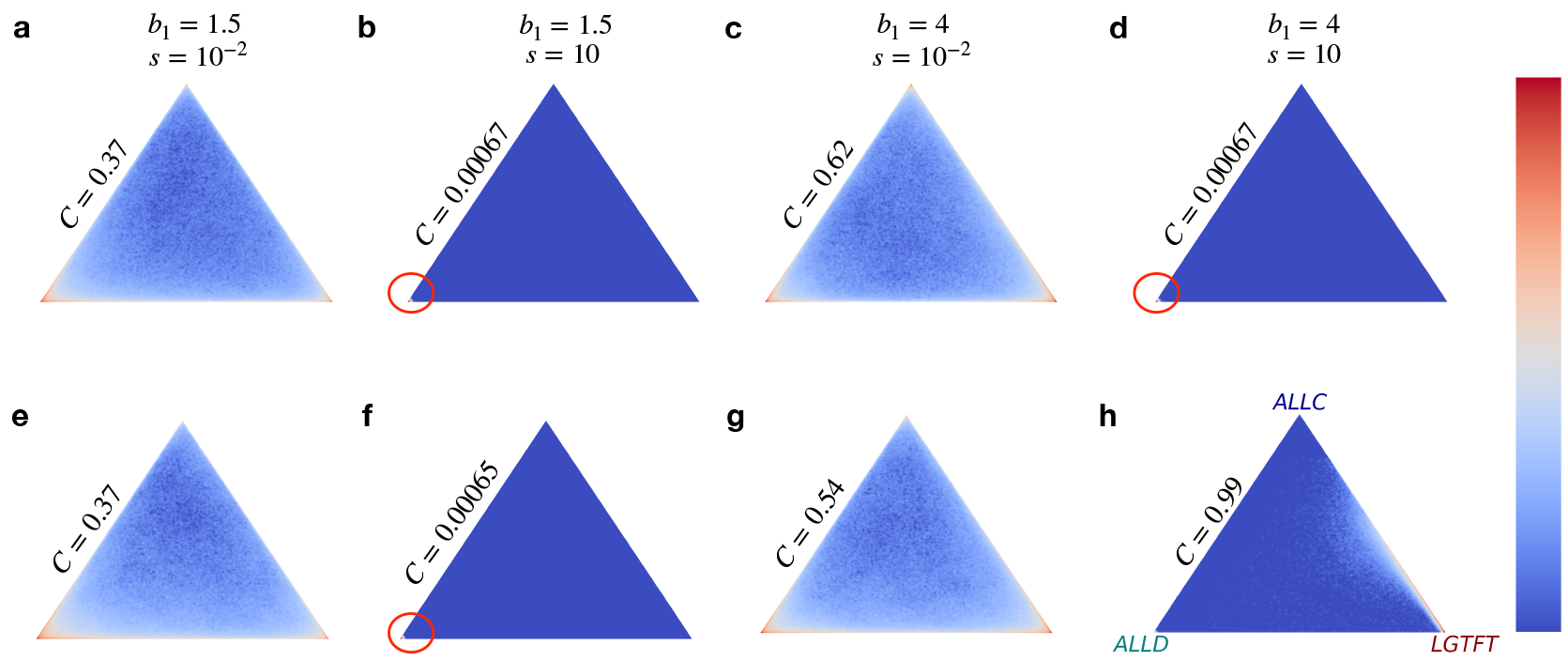
Heat map showing likelihood of different population compositions in the reduced strategy system {ALLC, ALLD, LGTFT}. Each point in the simplex represents the composition of the population averaged over the last 10^3^ time-steps out of a total simulation time of 10^5^ time-steps. The color of each point in the simplex denotes the number of times, out of 10^3^ independent simulations, the population appears with the specified abundance; with dark red being high and dark blue being low. (**a-d**) represent well-mixed and (**e-h**) represent structured population. The cooperation rate (*C*) is written above one edge of the simplex. Parameters: *N* = 200, *b*_2_ = 2, *c*_1_ = *c*_2_ = 1, *µ* = 0.001, *w* = 1. For LGTFT, *q* = 0.3.

For weak selection strengths, both LGTFT and ALLD are abundant in the population with ALLD being more dominant when *b*_1_ = 1.5 (Fig. 6.**a** and **e**) and LGTFT being more dominant when *b*_1_ = 4 (Fig. 6**c** and **g**) irrespective of the underlying population structure. This is also reflected in the cooperation rate which is larger in the latter case for both well-mixed (Fig. 6**c**) and structured populations (Fig. 6**g**). For large selection strengths, almost all players play the ALLD strategy in both well-mixed and structured populations when *b*_1_ = 1.5 as indicated by the red circle in (Fig. 6**b** and **f**). For *b*_1_ = 4, ALLD remains dominant in well-mixed populations (indicated by the red circle in Fig. 6**d**), while in structured populations, a mixed state of ALLC and LGTFT dominates, leading to a high cooperation rate (*C* = 0.99) in the population (Fig. 6**h**). These findings are consistent with the behaviour observed in our simulations carried out using the full strategy space at large *N* (Fig. 4d-4f), where the cooperation rates and abundance of full cooperation scenarios were found to be higher in multiplex-structured populations.

Our analysis with both an unconstrained strategy set as well as a reduced strategy set unambiguously confirm that a multiplex network structure facilitates the dominance of linked cooperative strategies (such as LGTFT) in multichannel games, resulting in high levels of cooperation in structured populations of all sizes. In contrast, selfish strategies like ALLD start dominating in well-mixed populations as the population size increases.

### 3.3 Evolution of cooperation in multiplex networks having identical topologies but distinct connections in each layer

So far, we have focused on multiplex networks with complete edge overlap between layers (𝒪= 1), which implies that each individual’s neighbors are identical across different social contexts. However, in real world scenarios, individuals are typically members of multiple social networks with each network having a distinct set of connections i.e. a distinct network structure. An individual may have some neighbours who are common across all layers of the multiplex and others that are unique to a specific layer of the multiplex. Since linking of strategies across layers is possible only for those neighbours of the focal player that are common across all layers of the multiplex, this effect can be investigated by systematically varying the number of common neighbours of each member of the population. We do so using a two-layer multiplex network, where we could adjust the structural similarities between the layers. Specifically, we considered two layers of Random regular network, each consisting of 100 nodes. By tuning the average edge overlap 𝒪 (eqn 2, for 2 layers 𝒪^*α,β*^ = 𝒪) of the multiplex, we can control the degree of similarity between the layers. Our model with variable edge overlap extends the work of [55] by providing a general framework to explore the impact of linking strategies on the evolution of cooperation for any fractions of common neighbors across layers. This perspective enhances our understanding of the effectiveness of strategy linking in more realistic scenarios represented by variable degrees of edge overlap of the multiplex network. Note that results in this and following sections have been obtained using the independent strategy update scheme only.

Fig. 7a shows the change in linked cooperation rate of population and Fig. 7b represents the change in cooperation rate of population as the fraction of common neighbors across layers is varied. As expected, the linked cooperation rate (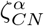, eqn. 11) calculated against common neighbors only, shows a monotonically increasing trend in both layers with increasing fraction of common neighbors. In contrast, the cooperation rate (*ζ*^*α*^, eqn 12), calculated against all neighbors (both common and unique) shows an U-shaped nature with change in fraction of common neighbors. Fig. 7b reveals that when individuals have a few common neighbours with whom strategies can be linked, strategy linking against common neighbors is not very effective in terms of boosting cooperation compared to the case without linking, i.e. when edge overlap 𝒪 = 0. Linking of strategies is detrimental to cooperation unless the degree of edge overlap is significantly high (i.e. 𝒪 ≳ 0.7).

**Figure 7:**
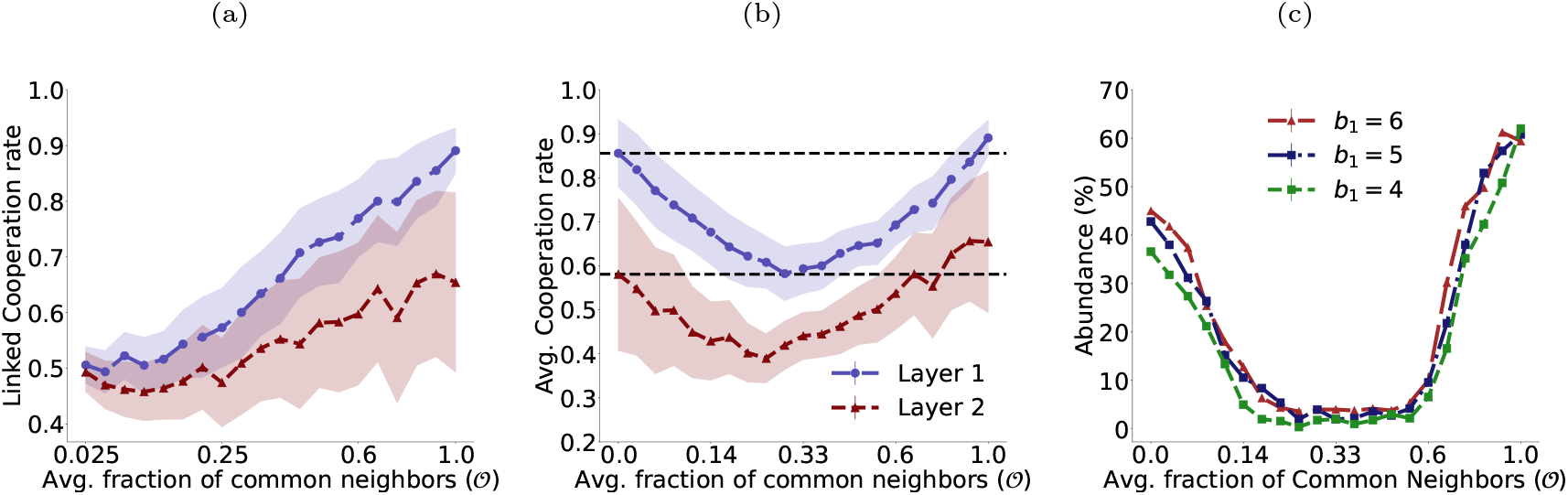
Impact of changing fraction of common neighbours in a multiplex network having RRN topology. **(a)** Change in the linked cooperation rate 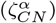, **(b)** cooperation rate (*ζ*^*α*^) in both layers, and **(c)** abundance of full cooperation scenarios; averaged over multiple independent simulations; with change in fraction of common neighbors across layers when an individual uses multi-game linked (unlinked) reactive strategies while interacting with common (unique) neighbours in both layers. Each point corresponds to the mean of 500 independent simulations each of which ran for 60,000 time-steps. The averaging over trials was done at the end of 60,000 time-steps. The shaded region depicts one sigma variation from the mean. The dashed horizontal lines in panel **(b)** represent the cooperation rate in each layer (1 & 2), when the fraction of common neighbors across layers 𝒪= 0 and are included as guides to the eye. Population size in each layer was *N* = 100, *k* = 20. All other parameters used are same as Fig.2.

When the benefit (*b*_1_) in layer 1 is very low (< *b*_2_), increasing the fraction of common neighbours has no effect on the linked cooperation rate (see Fig. 8a). However, for sufficiently large *b*_1_, increasing the fraction of common neighbours enhances the linked cooperation rate as expected since individuals with linked strategies are less prone to exploitation by selfish opponents in both games. On the other hand, the cooperation rate that considers both common and unique neighbours show an interesting behaviour. When the benefit (*b*_1_) in layer 1 is above a minimum threshold, increasing the fraction of common neighbours initially has a detrimental effect on the cooperation rate in layer 1 (see Fig. 8b). However, further sustained increase in the fraction of common neighbours eventually leads to an increases in the cooperation rate (see Fig. 8b). This behaviour generalizes the results depicted in Fig. 7b, where a U-shaped nature of the cooperation rate was observed for a fixed value of *b*_1_. An initial increase in the average fraction of common neighbours leads to a drop in cooperation rate in either or both layers of the multiplex as evident from the decrease in the abundance in full cooperation scenario (see Fig. 7c) where *ζ*^*α*^≥ 0.8 for *α* = 1, 2. For 𝒪*>* 0.6, the abundance of full cooperation scenario increases due to the increased advantage associated with strategy linking with a majority of neighbours, the cooperation rate also increases eventually reaching a maximum for complete overlap when 𝒪= 1. These results indicate that strategy linking is not always effective in enhancing the cooperation rate in multiplex structured populations and a significant amount of edge overlap between the two layers of the multiplex is necessary for strategy linking to yield the benefits of enhanced cooperation. To gain deeper insights into these results, we analyzed the strategies players employ against their common and unique neighbors as we varied the fraction of edge overlap (𝒪) between the two layers of the multiplex network (see Fig. 1 in SI). When edge overlap is zero or one, players on average use strategies similar to those observed in Fig. 2d,h, reciprocating altruistic acts with high probability. However, for intermediate edge overlap, we observed (see Fig. 1b,d in SI) that the evolved strategies on the average are less likely to reciprocate a co-player’s altruistic act in both games, leading to lower cooperation rates compared to 𝒪= 0 or 𝒪= 1 cases. This suggests that the degree of edge overlap can affect the nature of strategies that evolve with less altruistic strategies being employed against both common and unique neighbours for intermediate levels of edge overlap.

**Figure 8:**
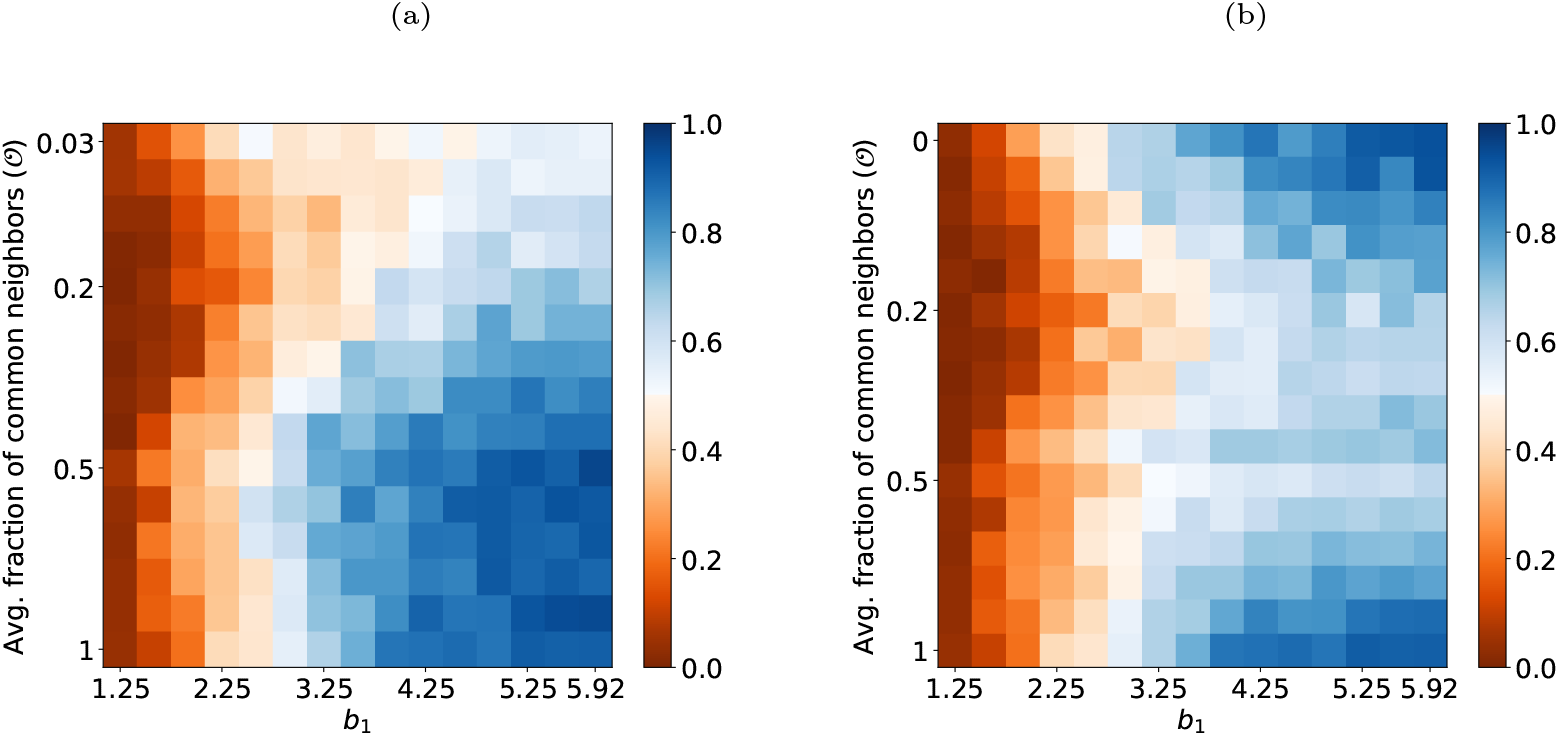
Impact of changing fraction of common neighbors and benefit of cooperation in layer 1. **(a)** linked cooperation rate 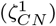 and **(b)** cooperation rate (*ζ*^1^) in layer 1. Each layer of the multiplex was created using RRN topology with degree *k* = 15. The value of each pixel corresponding to the respective types of cooperation rate was calculated at the end of the simulation that was run for 60,000 time-steps and averaged over 100 independent trials. The benefit in game 2 was kept fixed at *b*_2_ = 1.5 and *c*_1_ = *c*_2_ = 1. All other parameters used are same as in Fig. 7.

Fig.9 is a heat map showing the variation of the cooperation rate as the benefit of cooperation in both layers of the multiplex network is varied, for different degrees of edge overlap, corresponding to (a) 𝒪= 0, (b) 𝒪= 0.5, (c) 𝒪= 1. A comparison of the three panels indicate that the cooperation rate exceeds 0.5 over a larger region of *b*_1_-*b*_2_ space for zero and complete edge overlaps of the two layers of the multiplex. For intermediate degrees of edge-overlap (such as 𝒪= 0.5), increasing the benefits in layer *α* favours increased cooperative behaviour in that layer due to a higher chance of layer *α* cooperation scenarios emerging at the end of the simulation for each trial.

**Figure 9:**
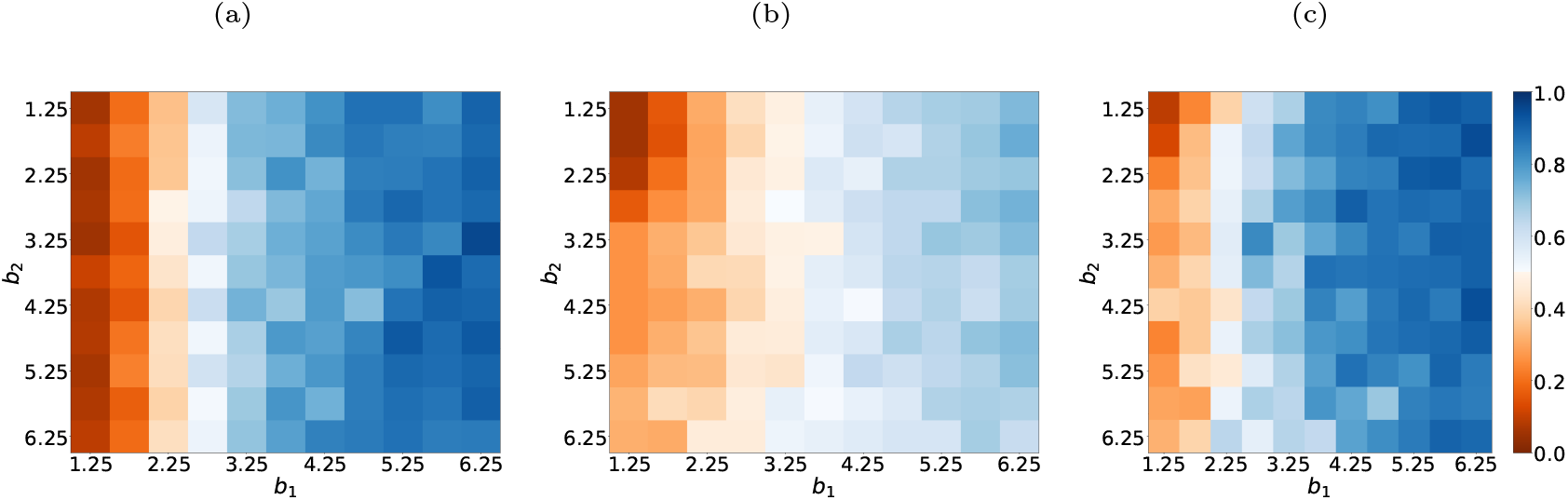
Heatmap depicting change in layer 1 cooperation rate with varying benefits associated with cooperation in both layers. The three panels correspond to cases where the degree of edge overlap is **(a)** 𝒪= 0, **(b)** 𝒪= 0.5, **(c)** 𝒪= 1. The multiplex network has RRN topology in each layer. Other parameters are same as those used in Fig. 7.

### 3.4 Incorporating imperfect memory as a cognitive constraint in repeated games

The success of conditionally cooperative strategies, such as GTFT in single repeated PD games and LGTFT in multiple repeated PD games depend on players having perfect memory of co-player’s past actions and correctly implementing their intended actions in the current round. While the impact of implementation error in repeated games is well-studied [6, 7, 59, 60, 61], most of these studies assume perfect memory for all players. Everyday experience and prior experiments [62],[63] indicate that people involved in interactions across multiple contexts often confuse past outcomes [64] and misremember previous actions. Previous theoretical studies on the evolution of cooperation have ignored the cognitive constraints of agents in social interactions. In the following, we study the impact of imperfect memory on the evolution of cooperation in multiple repeated games on multiplex network. For detailed derivations and discussions, see Sec.3.5 of SI.

To account for imperfect memory, we assume players misremember game 1 and game 2 outcomes with probabilities 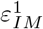 and 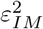, respectively. If a player is subject to imperfect memory in game 1, they will recall the opponent’s previous cooperation (*C*) as defection (*D*) and vice versa in game 1, while correctly recalling the co-player’s action (*a*_2_ ∈{*C, D*}) in game 2. If an error occurs in both games, a co-player’s cooperation in both games (*CC*) will be misremembered as defection (*DD*) in both games. In the former case of error in game 1 only, a player cooperates with probability 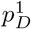 instead of 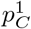 in game 1 against a unique neighbor in layer 1. For a common neighbor across layers, the player cooperates with 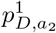 in game 1 and 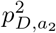 in game 2, instead of 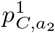 and 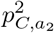, respectively. In the latter case of error in both games, a player cooperates with probabilities 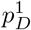 and 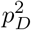 in game 1 and game 2, respectively, instead of 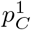 and 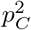 against unique neighbors in layers 1 and 2. Against common neighbors, cooperation probabilities shift to 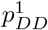 and 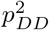 in game 1 and game 2, respectively, from 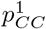 and 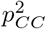. Here, we aim to understand the effects of imperfect memory in two cases: **(i)** when individuals use unlinked strategies against all neighbors (𝒪= 0), and **(ii)** when individuals use linked strategies against all neighbors across all layers (𝒪= 1). For unique neighbors, an error in one game does not affect the other, as each game is treated independently. In contrast, for common neighbors, errors in one game can significantly impact the other due to strategy linking across games. In line with this intuition, we varied error probability 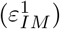 in game 1, while keeping perfect memory for game 2 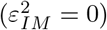. Fig. 10a shows that for unlinked strategies (𝒪= 0), the cooperation rate in game 2 remains around 85%, while in game 1, it declines rapidly from 94% to 0% with increasing error probability. Fig. 10b illustrates the variation in linked cooperation rates (𝒪= 1) in both games. Despite no errors in game 2, a moderate decrease in cooperation rate from 83% to 61% is observed due to the linking of strategies. Cooperation rate in game 1 becomes vanishingly small with increasing error probability. However, unlinked cooperation rate in game 1 decreases faster than the linked cooperation rate. To understand how errors in both games impact cooperation, we assumed error probability in both games are equal 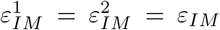 and we varied *ε*_*IM*_ continuously for both cases of 𝒪= 0 (Fig. 10c) and 1 (Fig. 10d). Consistent with previous experiments, we have seen that such errors have a significant negative impact on cooperation for scenarios with only unique and only common neighbors. For moderately high error probability (≳ 0.3) cooperation quickly diminishes in repeated games. Our results with imperfect memory underscores the importance of incorporating cognitive constraints in future research on repeated games.

**Figure 10:**
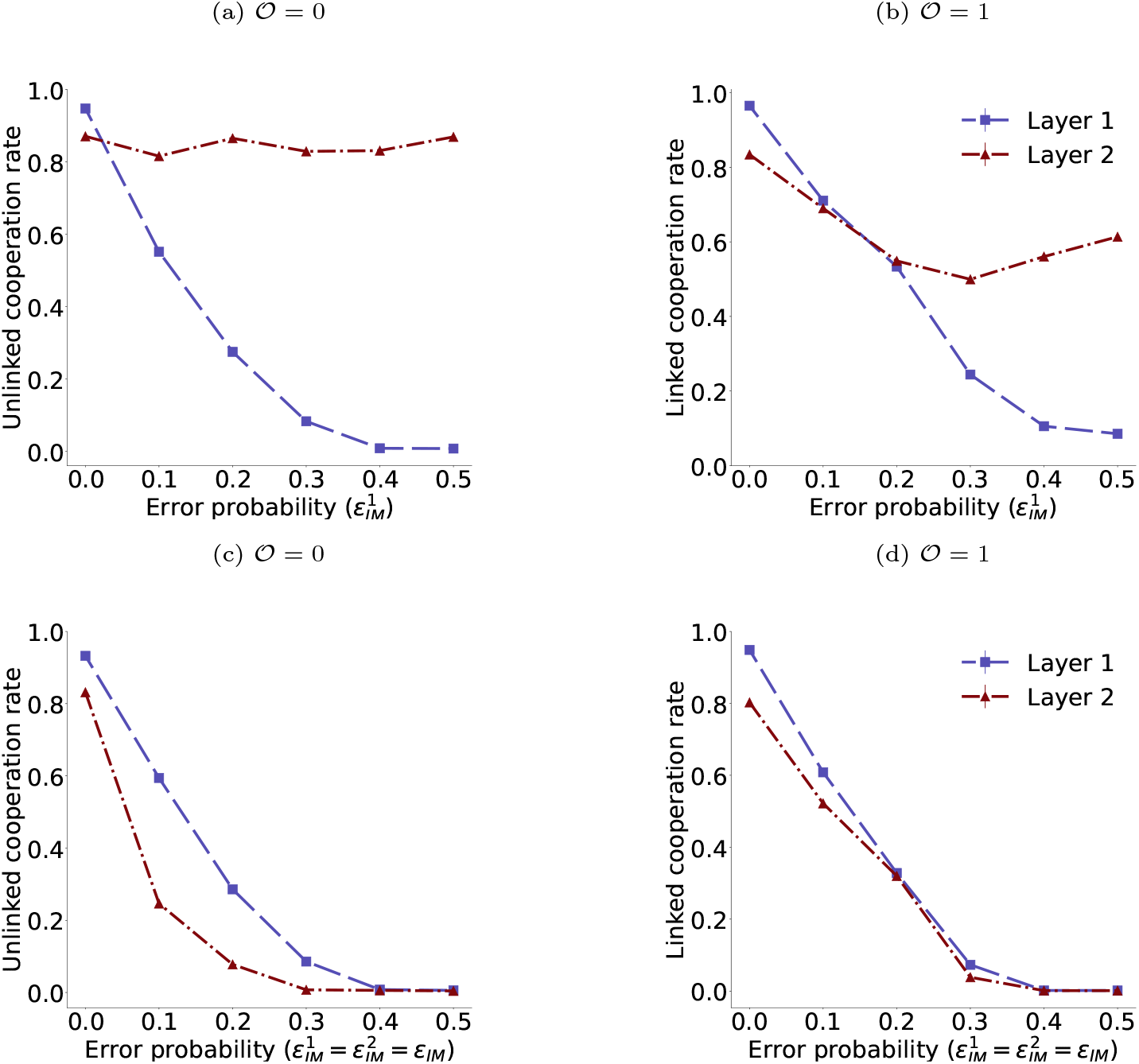
Impact of imperfect memory. The population’s cooperation rate when all individuals use either **(a, c)** unlinked strategy (𝒪= 0) or **(b, d)** linked strategy (𝒪= 1). In panels **a-b**, we varied the perception error in game 1 while game 2 was assumed to be error-free. In panels **c-d**, we varied the error in both games simultaneously. Incorporation of imperfect memory shows the negative impact of linking, as errors in game 1 reduce cooperation in game 2 (**b**), whereas unlinked cooperation in game 2 remains unaffected by increasing errors in game 1 (**a**). However, as errors increase in both games, cooperation diminishes rapidly for both unlinked (**c**) and linked (**d**) strategies. Each point represents the population’s cooperation rate, averaged over the last 1,000 time steps and 100 independent realizations. Each simulation of evolutionary dynamics ran for 10^5^ time-steps of independent strategy update. Both layers of multiplex network follow an RRN topology with nodes *N* = 100 and a degree, *k* = 4. Other parameters are the same as in figure 2.

### 3.5 Empirical multiplex network

We examined two real-world communities each interacting in multiple social contexts. We analyzed a two-layered, undirected multiplex network representing social (meeting for lunch) and working (research collaboration) relationships within the Aarhus University CS department [65], as well as business relationships and marriage alliances among Florentine families in the Renaissance [66]. The population size of each network ranged from *N* = 16 to *N* = 61. Across these communities, the average fraction of edge overlap varied from 𝒪= 0.296 to 𝒪= 0.34. Such networks deviate from the RRN networks studied in earlier sections since distinct nodes in each layer can have different degrees and different levels of edge overlap with corresponding nodes in the other layer. Fig. 11a, 11b shows visualisations of such multiplex networks. In our analysis, we assumed that if an individual shares a common interacting partner across all contexts (online/offline interactions, business partnerships/marriage ties), they will employ a linked strategy when interacting with that partner. We wish to understand how cooperation evolves in these real-world multiplex networks where individuals repeatedly engage in several interactions with various partners across multiple contexts.

**Figure 11:**
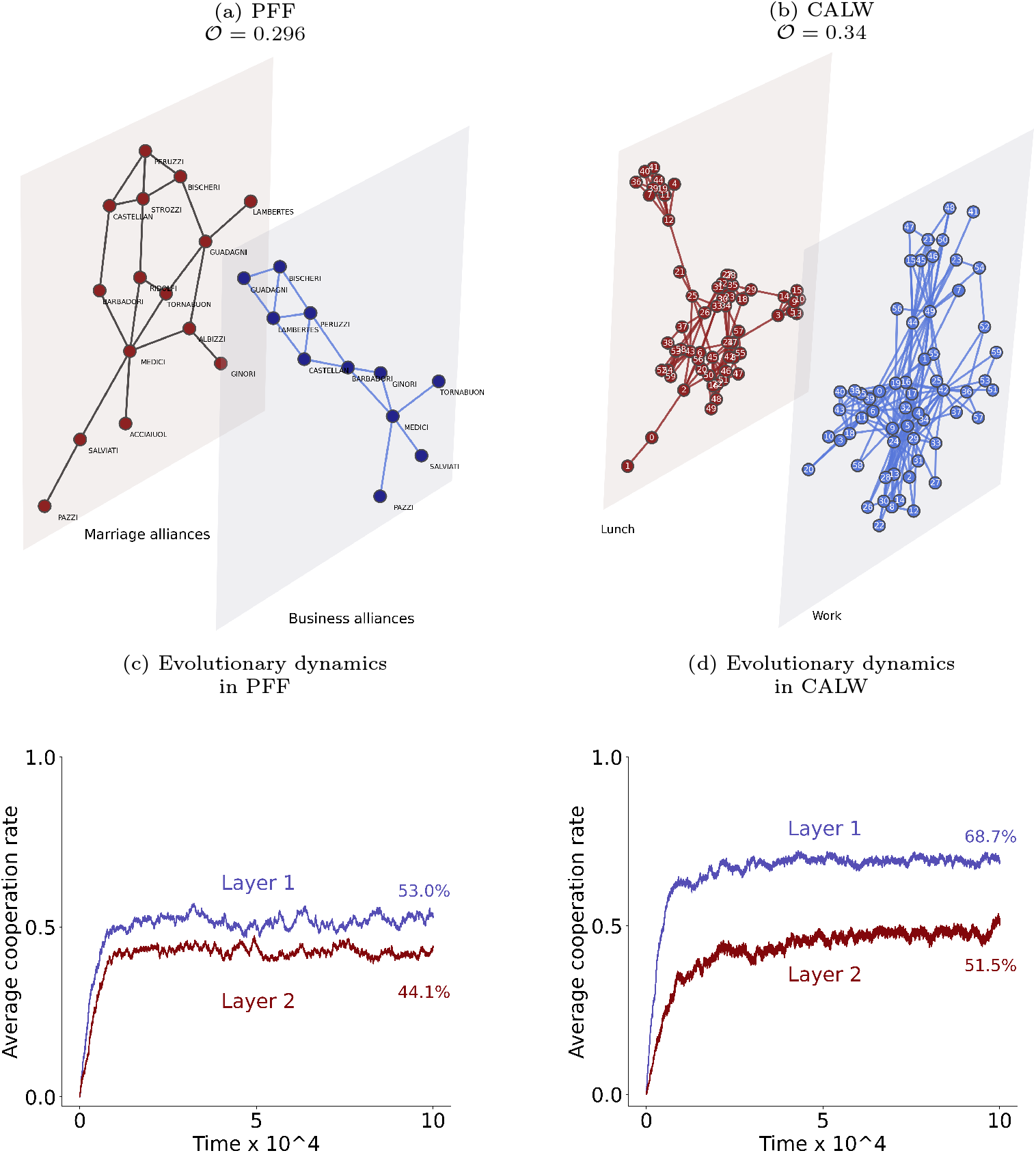
Evolution of cooperation in two-layered, undirected real-world multiplex networks: Visualizations of **(a)** the marriage alliance and business relationships of *N* = 16 Florentine families during the Renaissance (PFF) as a 2-layer multiplex network structure of Florentine families; **(b)** different offline relationships (eating lunch together and work-related collaboration) between *N* = 61 employees of the CS department in the Aarhus University (CALW). **(c) & (d)** shows the temporal evolution of populations’ cooperation rate in each layer. Each point is an average of 100 independent simulations using independent strategy update. One layer (blue) has a higher benefit of cooperation than the other (brown). Parameters used are the same as in Fig. 2.

We randomly selected a layer of the multiplex network and assumed the benefit of cooperation in this layer (blue layer, Fig. 11) is higher than in the other layer (brown layer). As expected, a higher level of cooperation evolved in the layer with greater benefit. Fig. 11c, 11d illustrates the temporal evolution of cooperation in two layers of the multiplex network of Florentine families and CS-Aarhus employees who share lunch and also collaborate on common work projects respectively. We observed that higher cooperation levels in both layers evolved in multiplex networks where the average edge overlap (𝒪) is higher. This finding aligns with our results obtained in Sec. 3.3 using a RRN topology.

## 4. Discussion

It is necessary to account for complex social interactions, that can occur across multiple domains, while attempting to decipher the underlying mechanisms that promote and sustain cooperation in human societies. It is in this spirit that we extended the framework of multichannel games to multiplex network. Moreover, such a framework allows us to understand how a strategic decision in one layer of the multiplex can affect decision in another layer and how strategy linking across layers of a multiplex affect the spread of cooperation in different layers. The efficacy of strategy linking in a multiplex network-structured population depends on a wide variety of factors like population size, degree of each network layer, extent of overlap between different network layers, perception errors and the strategy update rule. When the topology as well as the structure of the different layers of the multiplex are identical, strategy linking leads to greater abundance of the full cooperation scenario and higher cooperation rate in all layers of the multiplex compared to the mixed-population case, especially for large population sizes when the strategy of all layers are simultaneously updated. Moreover, strategy linking is found to induce high cooperation rates (60% or higher) in *both* layers only for large structural overlaps (𝒪 ≥ 60%) between the two layers. When the number of common neighbours is small, the benefits of linking strategies across layers do not accrue as much. The lack of coherence between linked and unlinked strategies ensures that the cooperation rate in both layers decreases with the initial increases in the fraction of common neighbours. However, when the fraction of common neighbours crosses a certain threshold (= 0.6), the ability of individuals to use linked strategies while playing with most of their neighbours enhance the cooperation rate in both layers. This is also manifest through the increase in abundance of full cooperation scenarios. Linking strategies also makes the system more susceptible to the negative consequences of errors in decisionmaking. Perception errors in just one game can lower the cooperation rate not only in that game but also in the game where there are no such errors.

Our work also opens up the possibility of using agent-based models to explore different types of strategy linking mechanisms across the layers of the multiplex network and their consequences on the evolution of cooperation. For instance, individuals may be more likely to cooperate in one layer if they are present in a cooperative environment in another layer. This underscores the importance of the strategy environment [67] in one layer affecting cooperative behaviour in other layers of the multiplex. Moreover, the issue of how the multiplex network structure affects the spread of cooperative behaviour in other social dilemmas like the public goods game, remains a pertinent question that is also open to future exploration. Our work suggests that multiplex networks provide a rich tapestry for emergent patterns of cooperative behavior that would not be seen in well-mixed populations and single layered networks.

## Acknowledgement

AB acknowledges the support provided by the Kepler Computing facility, maintained by the Department of Physical Sciences, IISER Kolkata. S.S. acknowledges partial support provided by a MATRICS grant no. (MTR/2020/000446, 2020-2023), given by SERB, India.

## Authors’ contributions

A.B.: conceptualization, data curation, formal analysis, investigation, methodology, software, visualization, writing-original draft, writing-review and editing; S.S.: conceptualization, formal analysis, methodology, resources, supervision, writing-original draft, writing-review and editing.

