## Supplementary Information for "Evolution of cooperation in multichannel games on multiplex networks"

September 19, 2024

**5 Description of real-world social networks used in the main text** **13**

### 1 Supplementary Figures

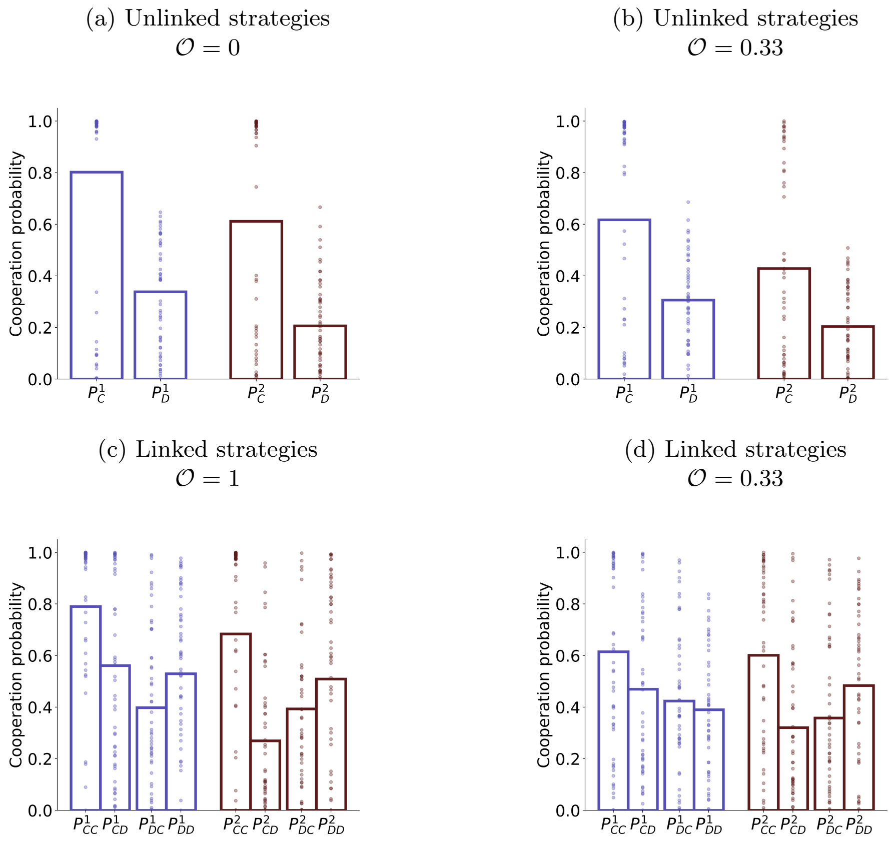

Figure 1: **Average strategy employed for different fraction of common neighbours:** Strategies players use (on an average) against their unique neighbors (panel **a-b**) and common neighbors (panel **c-d**) when the average fraction of common neighbors across two layers of the multiplex network are (**a**) 0, (**b, d**) 0.33, and (**c**) 1 respectively. $\mathcal{O} = 0.33$ indicates each individual has $O_i^{CN} = k/2 = 10$ common neighbors, where $k = 20$ is the degree of each node of the RRN in both layers of the multiplex network with $N = 100$ nodes per layer. Each bar represents the population's average of $p^{\alpha}_{a_1,a_2}(p^{\alpha}_{a_\alpha})$ against common (unique) neighbors in that respective layer, averaged over 100 independent realizations using independent strategy update, whereas dots represent 50 randomly sampled realizations of the simulation. Other parameter values used are the same as in Fig. 7 in the main text.

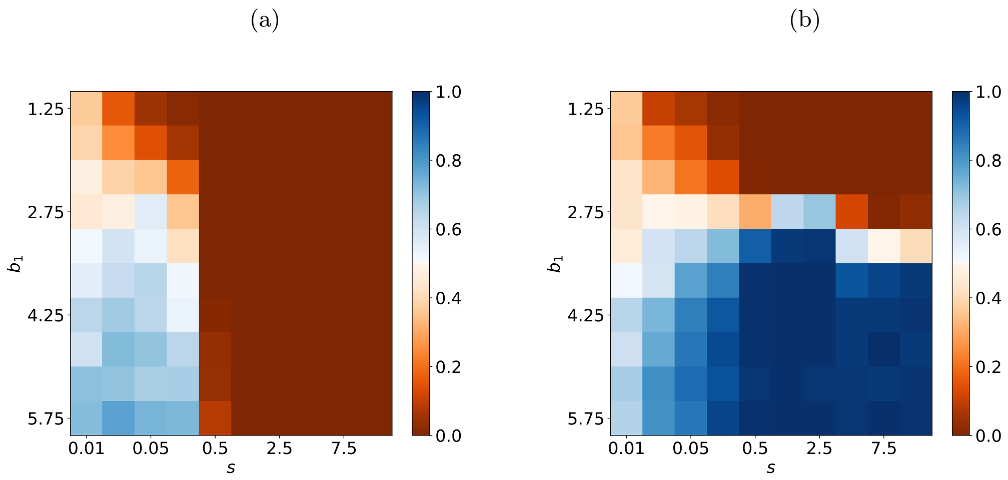

Figure 2: $b_1 - s$ **phase diagram using 3-strategy system.** **(a)** Well-mixed population with $k = N - 1$ and **(b)** Random regular network with $k = 4$. Each pixel corresponds to avg. cooperation rate which was averaged over the last $10^3$ time-steps and 100 independent simulations. Each simulation ran for $10^5$ time steps. Infinitely repeated games were considered in both cases with the benefit in game 2 fixed at $b_2 = 2$. Other parameter values used $c_1 = c_2 = 1, \mu = 0.001, N = 200$. For the LGTFT strategy, $q = 0.3$ was used.

### 2 Algorithm to create a multiplex network with desired edge overlap between layers:

To fine-tune $\mathcal{O}$ and create a multiplex with regular degree $k$ in both network layers, we initiated the process by creating two Random regular networks with same number of edges $k^{\alpha} = k^{\beta} = k' < k$, where $k'$ is initial degree of each layer, whose value was set as the desired number of common neighbors of each node ($O_i^{CN} = k' \;\; \forall \;\; i \in \{1, \ldots, N\}$). Hence we start with two identical network layers, where each node has degree $k'$ and the same set of edges in all the layers. Initially, the edge overlap is therefore maximal, $\mathcal{O}' = 1$. Subsequently, we increase the number of connections of all nodes in both layers in such a way that the number of common neighbors of each node remains unchanged. To grow both the network layers to achieve the desired degree $k$ and average fraction of common neighbors $\mathcal{O}$ for all layers, while maintaining an RRN structure, we carefully selected a set of valid edges for both networks by adhering to the following constraints. The new edges should not already exist within any of the two networks, the degree of each node in a particular layer should not exceed $k$, and any edge present in one network should not be included in the other (since a number of desired common neighbors ($k'$) has already been fixed in the initial configuration). From this set, connections were randomly established, ensuring the network retained its RRN characteristics with the average fraction of common neighbors given by $\mathcal{O}$.

### 3 Payoff calculation in multichannel games on a multiplex network

In this study, we have considered a multiplex network, where each individual can have distinct neighbors in different layers of the multiplex. In the multiplex, when an edge connecting two players, denoted as $i$ and $j$ respectively, exists across both layers of the multiplex network (common neighbor), player $i$ can conditionally choose to cooperate with player $j$ in game 1 of layer 1 as well as in game 2 of layer 2 based on the actions taken by player $j$ in *both* games during the previous round. For instance, if player $j$ defected in the previous round in layer 2, player $i$ can retaliate against $j$ in the next round by defecting in both games across both layers respectively. Conversely, in cases involving unique neighbors, such as the edge $i - k$ (see fig.1 of main text), player $i$ makes a decision regarding her action solely based on the action of player $k$ in the previous round in the specific game in the layer where the edge $i - k$ is present. We have assumed that each individual uses reactive strategies, wherein their decision to cooperate or defect is solely based on the actions of their opponents in the previous round only in *all* games. In a specific network layer $\alpha$, each individual is equipped with a linked reactive strategy given by $p^{\alpha}_{a_1, a_2, \ldots, a_m}$, where, $\alpha \in \{1, 2, \ldots, m\}$ for an $m$-layer multiplex network. This strategy represents the probability of cooperating in layer $\alpha$, given a tuple of actions $a := (a_1, a_2, \ldots, a_m)$, where $a_\alpha \in \{C, D\}$, represents the action taken by the opponent in the previous round in game $\alpha$. The initial round probability of cooperating against a common neighbor is given by $p^{\alpha}_{0 \ldots 0}$.
In the case of unique neighbors, where the edge $i - k$ has no edge overlap but exists only in layer $\alpha$, the strategy space against such neighbors becomes more restricted, specifically, the probability of cooperation for an $m$-layer multiplex becomes, $p^{\alpha}_{a_1, a_2, \ldots, a_m} = p^{\alpha}_{a_\alpha}$, where, $\alpha \in \{1, 2, \ldots, m\}$. The initial round probability of cooperating against a unique neighbor in layer $\alpha$ is $p^{\alpha}_0$.
The players' strategies are subject to implementation error with a small probability $\varepsilon$. With probability $\varepsilon$ players may occasionally misimplement their intended action. For error $\varepsilon > 0$, a players with strategy $\mathbf{p}$ effectively follows $(1 - \varepsilon)\mathbf{p} + \varepsilon(1 - \mathbf{p})$.

##### 3.1 Pairwise payoff against common neighbors in infinite repeated games

Infinitely repeated games involving two players $i-j$ utilizing the multi-game linked reactive strategies $\mathbf{p}$ and $\tilde{\mathbf{p}}$ respectively against their common neighbors, can be described using Markov chains. For two repeated donation games in two layers, the Markov chain can be defined on a state space with 16 possible states $((CCCC, CCCD, CCDC, \ldots, DDDD))$. In each of these states, the initial two letters signify the action of player 1, while the last two letters represent the action of player 2 in both game 1 and game 2 respectively, during the preceding round. For a more generalized version of a multiplex with $m$ layers where $m$ games are being played, Markov chain's current state is given by $\omega := (\mathbf{a}, \tilde{\mathbf{a}})$ and the next state is $\omega' := (\mathbf{a}', \tilde{\mathbf{a}}')$. Here, $\mathbf{a} = (a_1, a_2, \ldots, a_m) \in \{C, D\}^m$ is the action profile of first player and $\tilde{\mathbf{a}} = (\tilde{a_1}, \tilde{a_2}, \ldots, \tilde{a_m}) \in \{C, D\}^m$ is the action profile of second player in the current state. The transition probability[1] to move from the Markov chain's current state $\omega$ to the next state $\omega'$ is determined by the product of each player's individual probability to make the necessary decision in each game.

$$\Omega_{\omega,\omega'} = \prod_{\alpha=1}^{m} q^{\alpha}_{\omega,\omega'} \cdot \tilde{q}^{\alpha}_{\omega,\omega'} \tag{1}$$

Where individual probability is given as follows,

$$q^{\alpha}_{\omega,\omega'} = \begin{cases} p^{\alpha}_{\tilde{\mathbf{a}}} & \text{if } a'_\alpha = C \\ 1 - p^{\alpha}_{\tilde{\mathbf{a}}} & \text{if } a'_\alpha = D, \end{cases} \quad \text{and} \quad \tilde{q}^{\alpha}_{\omega,\omega'} = \begin{cases} \tilde{p}^{\alpha}_{\mathbf{a}} & \text{if } \tilde{a}'_\alpha = C \\ 1 - \tilde{p}^{\alpha}_{\mathbf{a}} & \text{if } \tilde{a}'_\alpha = D. \end{cases} \tag{2}$$

The $2^{2m} \times 2^{2m}$ transition matrix can be obtained by collecting all products $W = (\Omega_{\omega,\omega'})$. Since each probability term is between 0 and 1, every entry of $W$ is strictly positive. Thus, for infinitely repeated games from Perron-Frobenius theorem, it follows there exists a unique invariant distribution $\mathbf{v} = (v_\omega)$ which can be found by solving the linear equation $\mathbf{v} = \mathbf{v}W$. Each entry of this invariant distribution gives the probability to observe the outcome $\omega = (\mathbf{a}, \tilde{\mathbf{a}})$ over all rounds of multichannel games. Using this invariant distribution one can calculate the marginal distribution for each of the $m$ games in $m$ layers by computing,

$$v^{\alpha}_{a,a'} = \sum_{\omega} v_\omega e^{\alpha}_{\omega}(a', \tilde{a}') \tag{3}$$

$e^{k}_{\omega}(a', \tilde{a}')$ is an indicator function, whose value is one only when $a_k = a'$ and $\tilde{a_k} = \tilde{a'}$, otherwise zero. The marginal probability for each game $\alpha$ in layer $\alpha$ to be in one of the four possible states can be written as a vector $\vec{\mathbf{v}^{\alpha}} = (v^{\alpha}_{CC}, v^{\alpha}_{CD}, v^{\alpha}_{DC}, v^{\alpha}_{DD})$, where each term in the vector represents equilibrium frequency of the possible action of player 1 and player 2 respectively in layer $\alpha$. Player $i$'s(or $j$'s) repeated game expected payoff when playing against player $j$(or $i$) in each layer $\alpha$ can be calculated using this marginal distribution as follows,

$$\begin{aligned} \pi^{\alpha}_{ij} &= v^{\alpha}_{CC} R_\alpha + v^{\alpha}_{CD} S_\alpha + v^{\alpha}_{DC} T_\alpha + v^{\alpha}_{DD} P_\alpha \\ \pi^{\alpha}_{ji} &= v^{\alpha}_{CC} R_\alpha + v^{\alpha}_{CD} T_\alpha + v^{\alpha}_{DC} S_\alpha + v^{\alpha}_{DD} P_\alpha \end{aligned} \tag{4}$$

Where $\pi^{\alpha}_{ij}$ is player $i$'s repeated game pairwise payoff against common neighbor $j$ in layer $\alpha$. Player $i$'s (or alternatively, player $j$'s) pairwise average cooperation rate when playing against player $j$ (or alternatively, player $i$) in layer $\alpha$ respectively is

$$\begin{aligned} \gamma^{\alpha}_{ij} &= v^{\alpha}_{CC} + v^{\alpha}_{CD} \\ \gamma^{\alpha}_{ji} &= v^{\alpha}_{CC} + v^{\alpha}_{DC} \end{aligned} \tag{5}$$

##### 3.2 Pairwise payoff against common neighbors in finite repeated games

For finitely repeated games when two multigame linked reactive strategy players interact, their payoff and cooperation rate can be calculated in the similar method as described above. Suppose, the two players use the strategy $\mathbf{p}$ and $\tilde{\mathbf{p}}$ respectively. Then, the initial round probability to be on Markov chain's initial state $\omega := (\mathbf{a}, \tilde{\mathbf{a}})$ is given by,

$$v_\omega(0) = \prod_{\alpha=1}^{m} Q^\alpha_{\mathbf{a}} \cdot \tilde{Q}^\alpha_{\tilde{\mathbf{a}}} \tag{6}$$

Individual probabilities are calculated as follows,

$$Q^\alpha_{\mathbf{a}} = \begin{cases} p^\alpha_{0\ldots0} & \text{if } a_\alpha = C \\ 1 - p^\alpha_{0\ldots0} & \text{if } a_\alpha = D, \end{cases} \quad \text{and} \quad \tilde{Q}^\alpha_{\tilde{\mathbf{a}}} = \begin{cases} \tilde{p}^\alpha_{0\ldots0} & \text{if } \tilde{a}_\alpha = C \\ 1 - \tilde{p}^\alpha_{0\ldots0} & \text{if } \tilde{a}_\alpha = D. \end{cases} \tag{7}$$

The initial round outcome distribution is obtained by collecting all products $\mathbf{v}_0 = (v_w(0))$. Given the initial round distribution, all subsequent time distribution can be iteratively computed,

$$\mathbf{v}(t) = \mathbf{v}_0 W^t \tag{8}$$

Where $W$ is the standard transition matrix calculated using eqn. 1 and 2. The average distribution $\mathbf{v}$ obtained as follows [2],

$$\mathbf{v} = (1-w)\sum_{t=0}^{\infty} w^t \mathbf{v}(t) = (1-w)\mathbf{v}_0 \sum_{t=0}^{\infty}(wW)^t = (1-w)\mathbf{v}_0(I - wW)^{-1} \tag{9}$$

Here, $w$ is the continuation probability of another round of the game and $I$ is a $2^{2m} \times 2^{2m}$ identity matrix. $(I - wW)^{-1}$ is inverse of the respective matrix. Based on $\mathbf{v}$, pairwise payoff and cooperation rate for finitely repeated games can be calculated using eqn. 3, 4, and 5 in a similar way as infinitely repeated games described above.

##### 3.3 Pairwise payoff against unique neighbors

Each individual participates in multiple repeated games within a multiplex network, with some neighbors unique to each layer. Against these unique neighbors, games are treated independently. For repeated games in layer $\alpha$, a player's strategy takes the form $p^\alpha_{\tilde{a}_\alpha}$, where $a_\alpha \in \{C, D\}$ is a unique neighbor's action in game $\alpha$ in the previous round. This case is well-studied in the literature on repeated games.

Repeated games involving two players $i - k$ utilizing the unlinked reactive strategies $\mathbf{p}$ and $\tilde{\mathbf{p}}$ respectively against their unique neighbors, can be described as a Markov chain. For a multiplex network with $m$ layers, where $m$ independent games are played, the Markov chain's current state in layer $\alpha$ is given by, $\omega_\alpha{:=}(a_\alpha, \tilde{a}_\alpha)$ and the next state is $\omega'_\alpha{:=}(a'_\alpha, \tilde{a}'_\alpha)$. Here, $a_\alpha \in \{C, D\}$ is the action of the first player, and $\tilde{a}_\alpha \in \{C, D\}$ is the action of the second player in layer $\alpha$ in the current state. The transition probability of moving from Markov chain's current state $\omega_\alpha$ to the next state $\omega'_\alpha$ in layer $\alpha$ is written as,

$$\Omega^\alpha_{\omega_\alpha,\omega'_\alpha} = q^\alpha_{\omega_\alpha,\omega'_\alpha} \cdot \tilde{q}^\alpha_{\omega_\alpha,\omega'_\alpha} \tag{10}$$

Where individual probability is given as follows,

$$q^\alpha_{\omega_\alpha,\omega'_\alpha} = \begin{cases} p^\alpha_{\tilde{a}_\alpha} & \text{if } a'_\alpha = C \\ 1 - p^\alpha_{\tilde{a}_\alpha} & \text{if } a'_\alpha = D, \end{cases} \quad \text{and} \quad \tilde{q}^\alpha_{\omega_\alpha,\omega'_\alpha} = \begin{cases} \tilde{p}^\alpha_{a_\alpha} & \text{if } \tilde{a}'_\alpha = C \\ 1 - \tilde{p}^\alpha_{a_\alpha} & \text{if } \tilde{a}'_\alpha = D. \end{cases} \tag{11}$$

Collecting all products, the transition matrix for repeated games in layer $\alpha$ is given by, $W^\alpha = (\Omega^\alpha_{\omega_\alpha,\omega'_\alpha})$.
As the two players interact for infinitely many rounds, it converges to a unique invariant distribution $v^\alpha = (v^\alpha_{CC}, v^\alpha_{CD}, v^\alpha_{DC}, v^\alpha_{DD})$. Each entry $v^\alpha_{a\tilde{a}}$ represents the long-run probability of observing the game in the state $(a^\alpha, \tilde{a}^\alpha)$, where the first player chooses action $a^\alpha$ and the second player chooses action $\tilde{a}^\alpha$ in game $\alpha$. For $\mathbf{p}, \tilde{\mathbf{p}} \in (0,1)^2$, from Perron-Frobenius theorem the invariant distribution can be uniquely determined by solving the following eigenvector equation,

$$v^\alpha = v^\alpha W^\alpha \tag{12}$$

Using this invariant distribution and transition matrix, the payoff and cooperation rate for each layer $\alpha$ can be calculated (using equations 4 and 5) similarly to sections 3.1 and 3.2 for infinite and finite repeated games respectively.

##### 3.4 Total payoff in multiplex network

Player $i$'s accumulated payoff in layer $\alpha$ results from pairwise interactions with all her neighbors (both common and unique) in that layer. Thus, the total payoff to player $i$ in layer $\alpha$ is,

$$\begin{aligned}\pi_i^\alpha &= \frac{1}{k_i^\alpha}\left(\sum_{j\neq i}\delta_{1,o_{ij}}\pi_{ij}^\alpha + \sum_{j\neq i}(1-\delta_{1,o_{ij}})A_{ij}^\alpha\bar{\pi}_{ij}^\alpha\right)\\ &= \pi_i^{CN} + \pi_i^{UN}\end{aligned} \tag{13}$$

$\pi_{ij}^\alpha$ ($\bar{\pi}_{ij}^\alpha$) is pairwise payoff of $i$ against a common (unique) neighbor $j$ in layer $\alpha$ using strategy $p^\alpha_{a_1,a_2,\ldots,a_m}$ ($p^\alpha_{a_\alpha}$), where, $\alpha \in \{1, 2, \ldots, m\}$ respectively.
Player $i$'s total expected payoff from interactions across all layers is computed as,

$$\pi_i = \sum_{\alpha=1}^{m}\pi_i^\alpha \tag{14}$$

##### 3.5 Modelling cognitive constraints in repeated games

When players interact in repeated games, their behavior can be subject to cognitive constraints, such as imperfect recall of past actions, due to increased cognitive load associated with engaging in multiple parallel games. Such constraints can be incorporated into the framework of repeated and multichannel games as elaborated below.

###### 3.5.1 Imperfect memory

Consider a focal player interacting with a unique neighbor, in layer $\alpha$, who had cooperated in that layer in the previous round. With perfect memory, the focal player responds by cooperating with probability $p^\alpha_C$ in game $\alpha$. But with imperfect memory, the focal player may erroneously recall the previous action of her co-player in layer $\alpha$ as $D$ and as a consequence will respond by cooperating with probability $p^\alpha_D$. We call such errors perception errors.

Now consider a player interacting with a common neighbour who cooperated in game 1 and defected in game 2 in the previous round. With perfect memory, the focal player responds by cooperating with probabilities $p^1_{CD}$ in game 1 and $p^2_{CD}$ in game 2. However, with imperfect memory, if the erroneous recall happens, for example, in game 1 only, the focal player will misremember the previous actions of the co-player as $DD$. As a consequence, she will respond by erroneously cooperating with probabilities $p^1_{DD}$ in game 1 and $p^2_{DD}$ in game 2. If, on the other hand, erroneous recall happens in both games, the focal player will misremember the previous actions of the co-player as $DC$ and will respond by cooperating with probabilities $p^1_{DC}$ in game 1 and $p^2_{DC}$ in game 2.

Hence, in the case of common neighbours, imperfect recall in one game can affect actions in both games due to strategy linking. Conversely, for unique neighbors, errors in one game do not influence the other, as a result of interactions against different interacting partners in distinct layers. The following section gives the details of how imperfect memory is incorporated into our decision-making model for both unique and common neighbor scenarios.

*Unique neighbors:* With probability $\varepsilon^1_{IM}$ ($\varepsilon^2_{IM}$), players misremembers the action (regardless of whether it is $C$ or $D$) of the interacting partner in game 1 (game 2). These probabilities are *independent* of the nature of the co-player's last action ($a = C$ or $D$) in the respective games. When a player commits an error in layer $\alpha$, her original strategy $\mathbf{p}_{UN} = (p^\alpha_C, p^\alpha_D)$ changes to $\tilde{\mathbf{p}}_{UN} = (p^\alpha_D, p^\alpha_C)$ in the layer $\alpha$. Let $\mathbf{p}^\alpha_1$ and $\mathbf{p}^\alpha_2$ be the strategies against a unique neighbor for player 1 and player 2 respectively, in layer $\alpha$. We distinguish four cases corresponding to each game in layer $\alpha$ to show how strategies transform accordingly.

1. With probability $(1 - \varepsilon^\alpha_{IM})^2$ players commit no error and use their original strategy in game $\alpha$.

2. With probability $\varepsilon^\alpha_{IM}(1 - \varepsilon^\alpha_{IM})$, player 1 commits an error in game $\alpha$ but player 2 does not. In that case, player 1's original strategy in game $\alpha$, $\mathbf{p}^\alpha_1$ changes to $\tilde{\mathbf{p}}^\alpha_1$. Player 2 continues to use her original strategy in game $\alpha$.

3. With the same probability, $(1 - \varepsilon^\alpha_{IM})\varepsilon^\alpha_{IM}$, player 2 commits an error in game 1 but player 1 does not. In that case, player 2's original strategy in game $\alpha$, $\mathbf{p}^\alpha_2$ changes to $\tilde{\mathbf{p}}^\alpha_2$. Player 1 continues to use her original strategy in game $\alpha$.

4. With probability $(\varepsilon^\alpha_{IM})^2$ both player commits an error simultaneously in game $\alpha$. In that case, both of their strategy changes to $\tilde{\mathbf{p}}^\alpha_1$ and $\tilde{\mathbf{p}}^\alpha_2$ from $\mathbf{p}^\alpha_1$ and $\mathbf{p}^\alpha_2$ respectively.

The respective transition matrix for imperfect memory in game $\alpha$ can be written as,

$$\begin{aligned} W^\alpha_{IM} = (1-\varepsilon^\alpha_{IM})^2 W^\alpha(\mathbf{p}^\alpha_1, \mathbf{p}^\alpha_2) + \varepsilon^\alpha_{IM}(1-\varepsilon^\alpha_{IM})\Big(W^\alpha\left(\mathbf{p}^\alpha_1, \tilde{\mathbf{p}}^\alpha_2\right) \\ + W^\alpha\left(\tilde{\mathbf{p}}^\alpha_1, \mathbf{p}^\alpha_2\right)\Big) + (\varepsilon^\alpha_{IM})^2 W^\alpha(\tilde{\mathbf{p}}^\alpha_1, \tilde{\mathbf{p}}^\alpha_2) \end{aligned} \tag{15}$$

The transition matrix $W^\alpha(\mathbf{x}, \mathbf{y})$ on the right-hand side is defined as in the section 3.3. In the presence of imperfect recall error in any particular game, $W^\alpha_{IM}$ in equation 15 can be used to calculate the invariant distribution, payoff, and average cooperation rate of each player.

*Common neighbors:* Players misremember their co-players actions in game $\alpha$ from the previous round with probability $\varepsilon^\alpha_{IM}$ which is *independent* of the nature of the co-player's previous actions $(a, \tilde{a})$ in both games. For the two games considered, a player's original strategy against a common neighbor is represented as, $\mathbf{p}_{CN} = (p^1_{CC}, p^1_{CD}, p^1_{DC}, p^1_{DD}; p^2_{CC}, p^2_{CD}, p^2_{DC}, p^2_{DD})$. Based on this, let $\mathbf{p_1}$ and $\mathbf{p_2}$ be the common neighbor strategies of two players. We identify four distinct scenarios where a player misremembers her co-player's actions from the previous round in multichannel games, leading to a total of 16 possible misremembering cases for pairwise interactions between two players across two repeated games.

1. With probability $(1 - \varepsilon^1_{IM})(1 - \varepsilon^2_{IM})$, a player correctly remembers her co-player's action in both games and uses her original strategy $\mathbf{p}_{CN}$.

2. With probability $\varepsilon^1_{IM}(1 - \varepsilon^2_{IM})$, a player commits an error in remembering the co-player's previous action in game 1 but makes no error in game 2. In that case, the player's

original strategy changes in the following way,

$$\mathbf{p}_{CN} \rightarrow \mathbf{p}'_{CN} := (p^1_{DC}, p^1_{DD}, p^1_{CC}, p^1_{CD}; p^2_{DC}, p^2_{DD}, p^2_{CC}, p^2_{CD}) \tag{16}$$

3. With probability $(1 - \varepsilon^1_{IM})\varepsilon^2_{IM}$, a player commits an error in remembering the co-player's previous action in game 2 but makes no error in game 1. In that case, the player's original strategy changes in the following way,

$$\mathbf{p}_{CN} \rightarrow \mathbf{p}''_{CN} := (p^1_{CD}, p^1_{CC}, p^1_{DD}, p^1_{DC}; p^2_{CD}, p^2_{CC}, p^2_{DD}, p^2_{DC}) \tag{17}$$

4. With probability $\varepsilon^1_{IM}\varepsilon^2_{IM}$, a player commits an error in both games and misremember the co-player's previous round actions. In that case, the player's original strategy changes in the following way,

$$\mathbf{p}_{CN} \rightarrow \mathbf{p}'''_{CN} := (p^1_{DD}, p^1_{DC}, p^1_{CD}, p^1_{CC}; p^2_{DD}, p^2_{DC}, p^2_{CD}, p^2_{CC}) \tag{18}$$

The same four events may happen to the other player with the same probability. Using the same notation as before, the respective transition matrix becomes,

$$\begin{aligned}
W_{IM} = &(1 - \varepsilon^1_{IM})^2(1 - \varepsilon^2_{IM})^2 W(\mathbf{p_1}, \mathbf{p_2}) \\
&+ (1 - \varepsilon^1_{IM})^2(1 - \varepsilon^2_{IM})\varepsilon^2_{IM}\Big(W(\mathbf{p_1}, \mathbf{p''_2}) + W(\mathbf{p''_1}, \mathbf{p_2})\Big) \\
&+ \varepsilon^1_{IM}(1 - \varepsilon^1_{IM})(1 - \varepsilon^2_{IM})^2\Big(W(\mathbf{p_1}, \mathbf{p'_2}) + W(\mathbf{p'_1}, \mathbf{p_2})\Big) \\
&+ \varepsilon^1_{IM}\varepsilon^2_{IM}(1 - \varepsilon^1_{IM})(1 - \varepsilon^2_{IM})\Big(W(\mathbf{p_1}, \mathbf{p'''_2}) + W(\mathbf{p'_1}, \mathbf{p''_2}) \\
&+ W(\mathbf{p''_1}, \mathbf{p'_2}) + W(\mathbf{p'''_1}, \mathbf{p_2})\Big) + (1 - \varepsilon^1_{IM})^2(\varepsilon^2_{IM})^2 W(\mathbf{p''_1}, \mathbf{p''_2}) \\
&+ \varepsilon^1_{IM}(1 - \varepsilon^1_{IM})(\varepsilon^2_{IM})^2\Big(W(\mathbf{p''_1}, \mathbf{p'''_2}) + W(\mathbf{p'''_1}, \mathbf{p''_2})\Big) \\
&+ (\varepsilon^1_{IM})^2(1 - \varepsilon^2_{IM})^2 W(\mathbf{p'_1}, \mathbf{p'_2}) + (\varepsilon^1_{IM})^2\varepsilon^2_{IM}(1 - \varepsilon^2_{IM})\Big(W(\mathbf{p'_1}, \mathbf{p'''_2}) \\
&+ W(\mathbf{p'''_1}, \mathbf{p'_2})\Big) + (\varepsilon^1_{IM})^2(\varepsilon^2_{IM})^2 W(\mathbf{p'''_1}, \mathbf{p'''_2})
\end{aligned} \tag{19}$$

Given the transition matrix (equation 19), it is straightforward to calculate payoff and cooperation rates using equation 3, 4, 5.

### 4 Mathematical analysis using a reduced strategy set

In this section, we analytically study the evolutionary dynamics in well-mixed and structured populations under weak selection and rare mutation limits using a reduced strategy set. We assume that the multiplex network shares the same set of connections and topologies across layers. As a consequence, the two-layer network topology effectively reduces to a single-layer network topology. In this single layer, individuals concurrently play multiple games with their neighbors. This simplification allows us to obtain conditions for the LGTFT strategy to invade a resident population of purely selfish strategy players like ALLD (see section 4.1) in both well-mixed and network-structured populations. In Section 4.2, we analyze the 3-strategy set {ALLD, LGTFT, ALLC} subject to evolution via a selection-mutation process with a low mutation rate ($\mu \rightarrow 0$) and identify the analytical conditions under which cooperative strategies become abundant in the population in the weak selection ($s \rightarrow 0$) limit. We also carry out simulations to highlight the benefits of a network-structured population on promoting cooperation as $b_1$ and the selection strength ($s$) are varied.

#### 4.1 Analytical conditions for the invasion of ALLD by LGTFT

*Well-mixed population in the $N \to \infty$ limit*: For finitely repeated multichannel games between a pair of strategies $i - j$, payoff $\pi_{i,j}$ of strategy $i$ against $j$ can be calculated using the method described in section 3.2. For multichannel games between LGTFT and ALLD marginal probabilities (eqn. 9) for each game $\alpha \in \{1, 2\}$ can be written as

$$\begin{aligned} \mathrm{v}^1 &= (0, 1 - w + wq, 0, w - wq) \\ \mathrm{v}^2 &= (0, 1 - w + wq, 0, w - wq) \end{aligned} \tag{20}$$

Using these marginal probabilities, the pairwise payoff to each of the three strategies in each layer (given below) can be calculated using eqn. 4.

$$\begin{array}{c} \\ LGTFT \\ ALLD \\ ALLC \end{array} \begin{array}{c} \begin{array}{ccc} LGTFT & ALLD & ALLC \end{array} \\ \begin{pmatrix} (b_1 - c_1) + (b_2 - c_2) & -(c_1 + c_2)(1 - w + wq) & (b_1 - c_1) + (b_2 - c_2) \\ (b_1 + b_2)(1 - w + wq) & 0 & (b_1 + b_2) \\ (b_1 - c_1) + (b_2 - c_2) & -(c_1 + c_2) & (b_1 - c_1) + (b_2 - c_2) \end{pmatrix} \end{array} \tag{21}$$

Using the above payoff matrix 21, the evolutionary stability condition for the LGTFT strategy against ALLD in a *well-mixed* population is given by $\pi_{LGTFT,LGTFT} > \pi_{ALLD,LGTFT}$, which reduces to the condition

$$\frac{b_1 + b_2}{c_1 + c_2} > \frac{1}{w(1 - q)} \tag{22}$$

For infinitely repeated games ($w \to 1$), the above condition leads to,

$$q < 1 - \frac{c_1 + c_2}{b_1 + b_2} \tag{23}$$

The inequalities 22 and 23 reveal the benefits of linking strategies across multiple games on the success of cooperative strategies even in *well-mixed* populations. In the unlinked scenario, a TFT-like strategy ($q = 0$) can emerge in either layer if $\frac{b}{c} > \frac{1}{w}$ [3]. However, linking strategies across 2 games makes it easier for linked TFT ($LTFT$)-like strategies to be evolutionarily stable since the condition that needs to be satisfied $\frac{b_1+b_2}{c_1+c_2} > \frac{1}{w}$ is less stringent. This suggests that even if cooperation cannot emerge in the lower benefit game because ($\frac{b_2}{c_2} \not> \frac{1}{w}$), linking strategies across multiple games makes it easier for cooperation to be sustained since a less stringent condition $\frac{b_1+b_2}{c_1+c_2} > \frac{1}{w}$ needs to be satisfied.

In scenarios where $q > 0$, previous work [4] suggests that for a single well-mixed population, GTFT-like strategies ($(1, q)$) can be evolutionarily stable if $q < 1 - \frac{c}{b}$. However, linking of strategies across multiple games leads to a less stringent upper-bound on $q$ as given by the inequality 23; suggesting that LGTFT strategies can be more forgiving as long as strategies across both games are linked, leading to the LGTFT player cooperating with probability $q$ if her partner defects in even one of the games.

*Well-mixed population with finite size ($N$)*: In the case of a finite population, the success of LGTFT arises as a consequence of the condition that the fixation probability of the strategy must be larger than the neutral fixation probability ($1/N$) [5]. A strategy $i$ will be able to invade strategy $j$ in a finite but large population if

$$\pi_{i,i} + 2\pi_{i,j} > \pi_{j,i} + 2\pi_{j,j} \qquad \forall j \neq i \tag{24}$$

Using the payoff matrix 21 and the inequality 24, it follows that a single LGTFT player can take over a population of ALLD players if,

$$\frac{b_1 + b_2}{c_1 + c_2} > \frac{3 - 2(1 - q)w}{(1 - q)w} \tag{25}$$

The above condition 25 is also much easier to satisfy compared to the condition obtained for a single game.

*Structured population with finite but large N:* To understand the consequences of multiplex network structure on evolutionary game dynamics, we consider a Random regular network having a homogeneous degree ($k$) for all nodes. We adapt the methodology developed by Ohtsuki and Nowak [6] to obtain the condition for the success of the LGTFT strategy in such populations. According to [6], in a large regular network with finite degree $k$ the payoff between a pair of strategies $(i, j)$ changes from its mixed-population value in the following way:

$$\pi_{i,j} \to \pi_{i,j} + \frac{\pi_{i,i} + \pi_{i,j} - \pi_{j,i} - \pi_{j,j}}{k-2}, \qquad \forall i, j \tag{26}$$

The extra term on the right-hand side is due to local network structure and consequently limited interactions of each node in structured populations compared to mixed populations. Since the above inequality is derived in the large population limit, the contribution from the extra term vanishes when $k = N - 1$ thereby recovering the mixed population result. Note that, the network structure contribution also vanishes for diagonal terms and it is anti-symmetric. Thus the payoff matrix between the 3 strategies transforms to

$$\begin{array}{c} \\ LGTFT \\ ALLD \\ ALLC \end{array} \begin{array}{c} \begin{array}{ccc} LGTFT & ALLD & ALLC \end{array} \\ \begin{pmatrix} (b_1 - c_1) + (b_2 - c_2) & -(c_1 + c_2)(1 - w + wq) + H_1 & (b_1 - c_1) + (b_2 - c_2) + H_2 \\ (b_1 + b_2)(1 - w + wq) - H_1 & 0 & (b_1 + b_2) + H_3 \\ (b_1 - c_1) + (b_2 - c_2) - H_2 & -(c_1 + c_2) - H_3 & (b_1 - c_1) + (b_2 - c_2) \end{pmatrix} \end{array} \tag{27}$$

where, $H_1 = \frac{(b_1+b_2)(1-q)w-(c_1+c_2)(2-(1-q)w)}{k-2}$, $H_2 = 0$, and $H_3 = \frac{2(c_1+c_2)}{k-2}$.

Using the payoff matrix 27 in the inequality 24, the condition for $\rho_{LGTFT} > 1/N$, is found to be much less restrictive compared to mixed populations (inequality 25) in case of multichannel games and is given by

$$\frac{b_1 + b_2}{c_1 + c_2} > \frac{3 - 2(1-q)w}{(1-q)w} - \frac{3(1 - (1-q)w)}{(k+1)(1-q)w} \tag{28}$$

In the expression above, the critical benefit-to-cost ratio is an increasing function of $k$, thereby making it easier for LGTFT to invade ALLD in structured populations with lower degrees ($k << N - 1$) compared to well-mixed populations (where $k = N - 1$).

To validate our analytical predictions (inequality 28), we carried out simulations using a population comprising LGTFT and ALLD strategies (Fig. 5 in the main text). We generated an RRN structure consisting of $N = 100$ nodes. Initially, all nodes were set as ALLD, except for one randomly selected node, which was designated as a LGTFT player. In each iteration, we randomly chose an individual to update their strategy using a pairwise comparison rule. For each simulation run, we carried out these update steps until one of the two strategies got fixed in the population. To investigate how increasing the benefit in game 1 can facilitate the establishment of cooperation through the fixation of the LGTFT strategy, we estimated the fixation probability as a function of the benefit in game 1 while keeping the benefit in game 2 fixed at $b_2 = 2$. The cost of cooperation in both games was set to 1. We also compared the results for the multiplex network structured population with the mixed population scenario, represented by a complete graph.

The simulation results for the critical benefit-to-cost ratio agree well with our analytical predictions obtained above for structured populations (inequality 28) according to which $b_1 > 4.5, 5, 5.94$ for $k = 3, 5, N - 1$ respectively when $q = 0$ which corresponds to an LTFT) strategy (see Fig.5a of the main text). However, when $q = 0.3$ which corresponds to an LGTFT strategy, the predicted thresholds are $b_1 > 8.36, 9.28, 11.03$ for $k = 3, 5, N - 1$

respectively. For a complete graph ($k = N-1$), the predicted values are somewhat less than those found in our simulations since the theoretical assumption of $k << N$ necessary for deriving inequality 28 is no longer valid in this case. Nevertheless, these results indicate that multichannel games on network-structured populations can make it easier for cooperative strategies like LTFT and LGTFT to emerge, compared to well-mixed populations.

#### 4.2 Conditions for dominance in the 3-strategy system for weak and strong selection pressures

We first discuss the evolution of the 3-strategy system {ALLC,ALLD,LGTFT} subject to weak selection and low mutation pressures in a well-mixed population. Following the approach of Antal *et al.* [7], a strategy is identified as most favored by selection if it exhibits the highest frequency in the long-term average. In the remainder of this paper, we will refer to this long-term average frequency in the stationary state as "abundance".

Previous studies [7, 8] have shown that under weak selection (small $s$), each strategy $i$ in an $n$-strategy system has an approximate abundance of $1/n$, plus a deviation term $L_i$. For low mutation rates ($\mu \ll 1$), selection favors a strategy $i$ if:

$$L_i = \frac{1}{n}\sum_{j=1}^{n}(\pi_{i,i} + \pi_{i,j} - \pi_{j,i} - \pi_{j,j}) > 0 \tag{29}$$

Given the payoff matrix for the 3-strategy system 21 in a well-mixed population, the linear coefficients $L_i$ according to eqn. 29 can be simplified to,

$$\begin{aligned} L_{LGTFT} &= \frac{1}{3}[(1-q)(b_1+b_2) - (1+q)(c_1+c_2)] \\ L_{ALLD} &= \frac{1}{3}[-(1-q)(b_1+b_2) + (q+3)(c_1+c_2)] \\ L_{ALLC} &= -\frac{2}{3}(c_1+c_2) \end{aligned} \tag{30}$$

For low mutation rates and for infinitely repeated games ($w \to 1$), ALLC can never dominate for any values of $b_1$ and $b_2$. This is straightforward to understand since ALLC is neutral with respect to LGTFT and is exploited by ALLD in all games. With increasing benefit-to-cost ratios in different games, the relative abundance of these strategies changes with the nature (*i.e.* value of $q$) of the LGTFT strategy. According to eqn. 30, the ALLD strategy has an abundance greater than 1/3 if $q > 2q^* - 1$ while LGTFT strategy has an abundance greater than 1/3 if $q < q^*$, where $q^* = \frac{(b_1+b_2)-(c_1+c_2)}{(b_1+b_2)+(c_1+c_2)}$ . Since $q^* > 2q^* - 1$, for $2q^* - 1 < q < q^*$, both LGTFT and ALLD strategies have abundances greater than 1/3. The condition $q < 2q^* - 1$, which implies that only LGTFT dominates with a frequency greater than 1/3, can only be true if $2q^* - 1 \geq 0$ (since $q \geq 0$) which leads to the condition

$$\frac{b_1+b_2}{c_1+c_2} \geq 3 \tag{31}$$

Note that inequality 31 doesn't guarantee that the inequality $q < 2q^* - 1$ will be satisfied. Thus, inequality 31 is a necessary but not sufficient condition for only LGTFT to become the most abundant strategy in the population.

Similarly, we can calculate $L_i$ according to eqn. 29 for network structured population using the payoff matrix 27. However, the conditions for an abundance of different strategies in a structured population are exactly the same as in a well-mixed population obtained above. Thus, in the limit of weak selection, network structure offers no help in promoting cooperation in multichannel games compared to a well-mixed population. However, as the selection strength increases, the distinctions between evolutionary outcomes in well-mixed and network-structured populations become clearly manifest.

Fig. 2 shows the average cooperation rate as a function of benefit in game 1 ($b1$) and selection strength $s$ for well-mixed (Fig. 2a) and RRN structured (Fig. 2b) populations with degree $k = 4$. A well-mixed population is completely dominated by ALLD for any value of $b_1$ when selection strength crosses a certain threshold ($s > 0.5$). However, the outcome changes dramatically for an RRN structured population. When the benefit in game 1 is sufficiently large ($b_1 > 2.75$), the LGTFT strategy becomes the most abundant strategy even in the strong selection regime (Fig. 2b) in the network structured population, resulting in a high cooperation rate.

Multichannel games in a well-mixed but large population fail to sustain cooperation, even if strategies across games are linked. In contrast, we have shown that cooperation can be sustained across all population sizes when multiple games are played across different layers of a multiplex network-structured population.

### 5 Description of real-world social networks used in the main text

We investigate three empirical social networks (datasets available at `https://icon.colorado.edu/#!/`) in the main text. Some of these networks have more than two layers, but for simplicity, we focus on the evolution of cooperation in two layers. We either used two layers from the original network or combined multiple layers into two layers of interest.

1. Pedgett Florentine Families: The multiplex social network consists of 2 layers depicting marriage alliances and business relationships of Florentine families in the Renaissance. There are 16 nodes in total, each engaged in different social relationships with each other. The multiplex is undirected and unweighted.

2. CS-Aarhus: The multiplex social network comprises of five types of online and offline relationships (Facebook, Leisure, Work, Co-authorship, Lunch) among employees of the Computer Science department at Aarhus University. In this study, we created two different multiplex networks from the original five layers: one based on online (Facebook, Co-authorship) and offline (Lunch, Leisure, Work) relationships, and another based on employees who share lunch and also work together.

[2] Karl Sigmund. *The calculus of selfishness*. Princeton University Press, 2010.
